# Metagenomics and metabarcoding experimental choices and their impact on microbial community characterization in freshwater recirculating aquaculture systems

**DOI:** 10.1101/2022.07.05.498813

**Authors:** Jessica Rieder, Adamandia Kapopoulou, Claudia Bank, Irene Adrian-Kalchhauser

**Affiliations:** Institute for Fish and Wildlife Health, Department of Infectious Diseases and Pathobiology, Vetsuisse Faculty, University of Bern. Länggasstrasse 122, 3001 Bern, Switzerland; Division of Theoretical Ecology and Evolution, Institute of Ecology and Evolution, University of Bern. Baltzerstrasse 6, 3012 Bern, Switzerland; Swiss Institute of Bioinformatics, Quartier Sorge - Batiment Amphipole, 1015 Lausanne, Switzerland

**Keywords:** aquaculture, recirculating aquaculture system (RAS), microbial communities, DADA2, 16S rRNA gene, amplicon sequencing, shotgun metagenomics, ASVs, MiSeq, Pac-Bio, short- and long-reads

## Abstract

Recirculating aquaculture systems (RAS) heavily depend on microbial communities to maintain water quality. These communities therefore influence the growth, development, and welfare of farmed fish. With the increasing socio-economic role of fish farming e.g. regarding food security, an in-depth understanding of aquaculture microbial communities is also relevant from a management perspective. However, the data situation regarding the composition of microbial communities within RAS is patchy. Since this is partly ascribed to method choices, there clearly is a need for accurate, standardized, and user-friendly methods to study microbial communities in aquaculture systems.

Here, we compare the performance of 16S amplicon sequencing, Pac-Bio long-read amplicon sequencing, and amplification-free shotgun metagenomics in the characterization of microbial communities in two commercial-size RAS fish farms. We show that, even though primer choice affects read quality, diversity, and assigned taxa, distinct primer pairs uncover similar spatio-temporal patterns between sample types, farms, and time points. We find that long-read amplicons underperform regarding quantitative resolution of spatio-temporal patterns, but allow for species-level identification of functional services and pathogens. Finally, shotgun metagenomics data identified fungi, viruses, and bacteriophages, opening avenues for an exploration of natural approaches regarding antipathogenic treatments. Overall, the datasets agreed on major prokaryotic players.

In conclusion, different sequencing approaches yield overlapping and highly complementary results, with each contributing data no other approach could. Such a tiered approach therefore constitutes a practical and cost-effective strategy for obtaining the maximum amount of information on aquaculture microbial communities. These data could lead to better farm management practices and at the same time inform basic research on community evolution dynamics.

## Introduction

Recirculating aquaculture systems (RAS) are a valuable alternative to the limited sustainable capacity of capture fisheries. They are discussed as a long-term sustainable offset for capture fisheries [1] and as a means to meet the nutritional demand for high-quality animal protein. RAS cultivate freshwater species such as rainbow trout (*Oncorhynchus mykiss*), pike-perch (*Stizostedion lucioperca*), Arctic char (*Salvelinus alpinus*), and sturgeon (order *Acipenseriformes*) [2] and range from small privately-owned enterprises to industrial-sized corporations. Importantly, the in-door, closed-circuit design of RAS provides independence from seasonal conditions, allows for biosecurity measures, and reduces the product-to-market distance when situated inland [3].

Microbial communities in RAS play a crucial role in overall system success, nutrient cycling, water quality, and animal health [1], [4]–[12]. These communities are often actively maintained in the biofilter section of the system, which is designed to maximize the surface area with sand, granulated active carbon, or synthetic carrier material, and perform an array of services, such as removing toxic metabolic products (e.g., ammonia, nitrite, nitrate, sulfide, and sulfate), and organic waste. Prominent representatives of the oxidizing ammonia genera found in RAS are, for example, *Nitrosomonas*, *Nitrosospharea*, and *Nitrosospira* [13], as well as ammonia-oxidizing archaea and *Nitrotoga* species [14], [15].

Conversely, pathogenic components of microbial communities in RAS constitute a significant challenge for the fish farm industry. Fish-related disease outbreaks threaten the livelihood of farmers and food security [16] and incur an estimated $6 billion loss yearly [17] as a result of stock loss. Also, water-associated off-flavoring bacterial groups may adversely impact the quality of the final product [13]. Various management approaches, such as cleaning and disinfection, aim to reduce opportunistic pathogen species such as *Aeromonas* or *Flavobacterium* [18] but could potentially open niches for pathogenic species and promote undifferentiated microbial growth.

Accordingly, managing the microbial communities that maintain water quality in RAS poses complex challenges. It has been proposed that monitoring and targeted manipulation of RAS microbial communities, based on a thorough characterization of interactions and community dynamics, may improve aquaculture management strategies [19]–[21]. However, RAS microbial research lags behind compared to other microbe-dependent industries such as wastewater treatment. The interactions between different compartments, management operations, microbial community structure, and how community assemblages differ across facilities are only beginning to be understood [1]. Previous microbial studies have analyzed the biofilter communities in RAS farming lumpfish (*Cyclopterus lumpus L.*) [8], Atlantic salmon (*Salmo salar*), Pacific white shrimp (*Litopenaeus vannamei*), half-smooth tongue sole (*Cynoglossus semilaevis*) and turbot (*Scopthalmus maximus*) [22], but have not investigated other RAS compartments. Inter-study comparisons are not straight-forward because of non-standardized protocols (e.g., DNA extraction, amplification, sequencing platform, or taxonomic assignment), which impact the results and conclusion [23]–[27]. In addition, global studies are scarce [28], and as a result, the characterization of RAS microbial community patterns and keystone taxa remains incomplete.

This study investigates the effect of sampling and analysis strategies on the inference of microbial community composition in RAS. We collected data from two freshwater RAS, compared four primer sets, and generated and compared three types of sequencing data (Figure 1) to identify key microbial dynamics and improve future sampling and methods decisions. First, we show that primer-specific results at early analysis steps do not lead to distinct biological conclusions. Second, we demonstrate that 16S short-read sequencing is sufficient to detect spatio-temporal developments and dynamics in the context of a RAS system. Finally, we demonstrate and discuss the distinct value of short amplicon, long amplicon, and shotgun metagenomics approaches depending on the research question and use a combination of the different sequencing results to describe the spatio-temporal patterns and identify microbials in different compartments of RAS.

**Figure 1:**
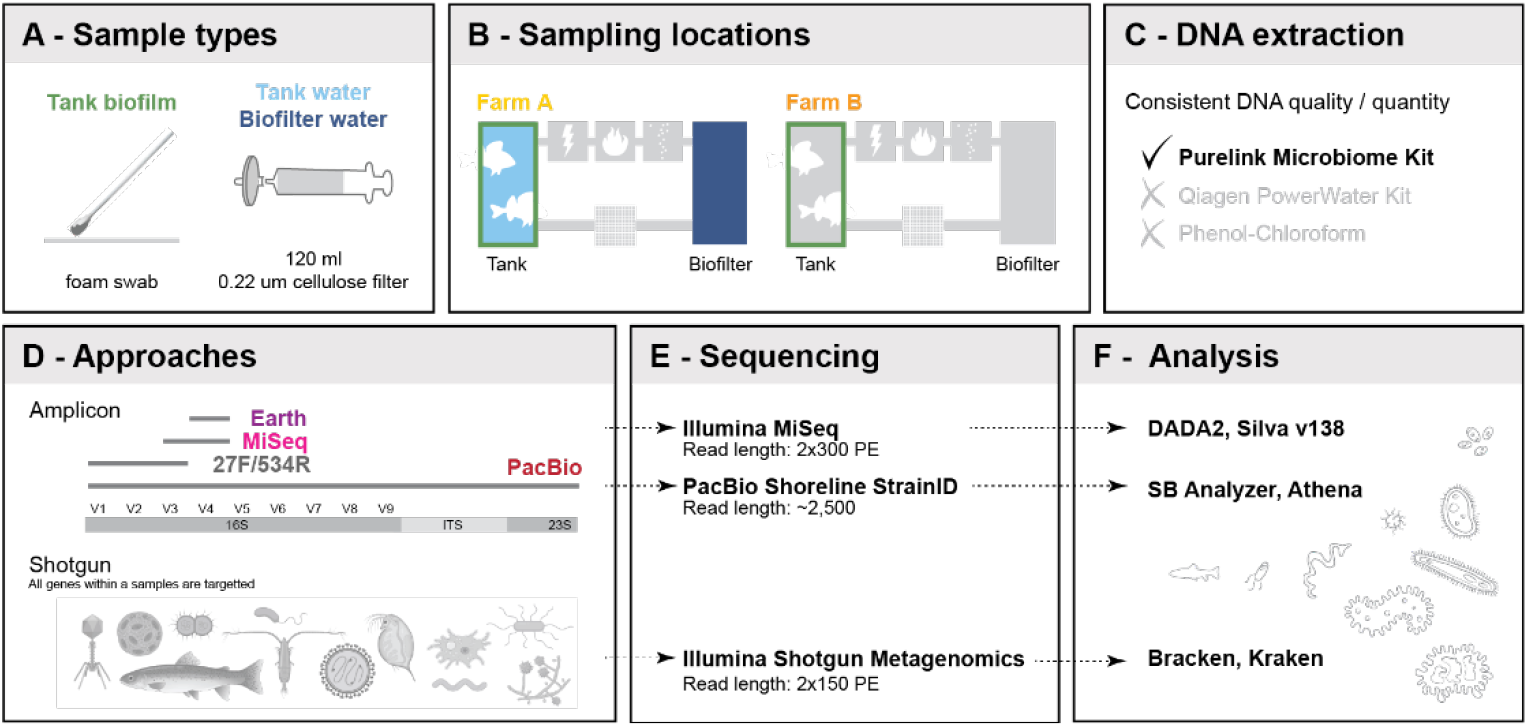
Study design and experimental steps. A: Three types of samples were taken. Tank biofilm was collected by rubbing a foam swab against the sidewall of a tank. Tank and biofilter water was collected by filtering 120ml of water through a 0.22 um cellulose filter. B: Samples were taken at two farms, A and B. In farm A, all three sample types were collected. In farm B, sampling focused on tank biofilm. C: Three DNA extraction methods were compared. The Purelink Microbiome Kit outperformed other DNA extraction methods in DNA quality, quantity, and consistent yield. D: Two types of amplicon approaches (short and long reads) and an amplification-free shotgun approach were used. E: Short amplicons were sequenced on an Illumina MiSeq with a v3 2×300PE kit. Long amplicons were sequenced with the Pac-Bio Shoreline StrainID kit, producing an average read length of 2,500bp. Shotgun sequencing was performed with a 2×150PE illumina kit. F: Illumina MiSeq sequencing data were processed with the DADA2 pipeline, and ASVs were blasted against the SILVA v138 database for taxonomic assignment. Pac-Bio long-read data was processed with the SBAnalyzer program and taxonomically assigned using Athena. Shotgun metagenomics sequencing data were processed with an in-house pipeline that uses the Kraken-Bracken method.

## Materials and Methods

### Sampling sites

The study includes two commercial-size Swiss RAS farms (A and B) with distinct ownership and operational management procedures. Farm A breeds perch (*Perca fluviatilis*), raising offspring from egg to approximately 15g and features several life-stage-specific circuits with independent filtration systems. Fish are moved to the next circuit when they reach a certain cutoff weight. After each move, a stringent disinfection regimen is applied. First, the biofilter is disconnected from the circuit to protect the microbial community from disinfection solutions. Next, the tanks are emptied, followed by a four-step cleaning regimen, 1) a high-power jet wash with hot water, 2) brushing down the tank walls and floor with soap, 3) a static acid-base treatment of the tanks and pipes with neutralizing steps in between, and 4) spraying the tanks with alcohol. The tanks are then dried entirely before refilling and restocking with the next batch of fish. Farm A uses multiple feed brands depending on the life stage of the fish (Bernaqua, BioMar, and Alltech Coppens).

Farm B is situated >100 km from Farm A in a different catchment. Farm B raises two fish species: perch, obtained from Farm A at around 15g, and pike-perch (*Sander lucioperca*), obtained at the fingerling stage from another provider. Both species are raised to slaughter weight within a single circuit in concrete tanks. Cleaning regimens are applied once a tank is emptied. However, because of grading and moving the fish into new tanks, which might already be occupied, there is no strict cleaning disinfection timeline. The disinfection protocol consists of emptying tanks, washing the tank with high-pressure hot water, spraying Virkon S as a disinfection solution, refilling with water, and then stocking with the next batch of fish. Farm B uses Alltech Coppens Supreme of varying pellet sizes according to fish size. Both farms use agitated biofilters with floating plastic biofilter carriers to supply the necessary surface area to foster microbial communities. Farm identities and locations are confidential.

### Sample types

Three sample types were collected: tank biofilm, tank water, and biofilter water (Figure 1A). First, biofilm samples were collected with a sterile, single-use foam swab (Merck – product was discontinued) by rubbing one side of the swab back and forth approximately ten times across a ~10×10 cm area of the tank wall about 6 cm below surface water level and repeating the procedure on the same area with the other side of the swab. After swabbing, the swab was placed into a 2 ml Eppendorf tube, the stick was broken off, and the closed 2 ml tube was placed on wet ice. Biofilm replicates were taken with an approximately 2 cm gap between them. Next, using a sterile 500 ml plastic beaker, we collected 500 ml of water from the same tank where the tank biofilm sample was collected, followed by on-site filtering of 120 ml of water using a 60 ml sterile, single-use syringe (Faust) and a 0.22 um mixed cellulose filter (Millipore, Merck) contained in a Whatman 47 mm plastic filter holder (Whatman, Merck). Replicates were taken from the same beaker, thoroughly mixing the water before the replicate was taken. After filtration, the filters were placed in a 2ml Eppendorf tube and stored on ice. Finally, biofilter water samples were collected, with a new sterile beaker, in the same way and from the same circuit as tank water and biofilm samples. All samples were transported back to the Institute for Fish and Wildlife Health, University Bern, on wet ice and stored at −80°C until further processing.

### Sampling scheme

Sampling aimed to maximize insights into differences and similarities between replicates, sample types, time points, analysis methods, within-farm compartments, and farms. In Farm A, two sampling events on different dates occurred in the H circuit that houses 12-15g perch. The first sampling event took place on June 25^th^, 2020 and consisted of collecting tank wall biofilm (samples 4-6), tank water from the same tank as the biofilm (samples 7-9), and biofilter water (samples 10-12). This sampling took place less than a week after the last cleaning of the tanks. A second sampling took place on November 4^th^, 2020 and involved the collection of tank wall biofilm (samples 1-3), from a second tank within the H circuit, several weeks after the last cleaning of the tank. In farm B, sampling took place on November 23^rd^, 2020, that consisted of collecting tank wall biofilm from two tanks (samples 13-15 (tank 1) and 16-18 (tank 2), within the main circuit, E. Two negative control samples were taken at the June 25^th^, 2020, sampling event but not sequenced. An overview of all samples is provided in Additional file 1. The negative water control was filtered the same way as the on-site water samples, using distilled water instead of system water. The negative swab sample consisted of unpacking a swab on-site and placing it into a 2 ml tube without swabbing a surface.

### DNA extraction

Three DNA extraction methods were tested on pre-trial water and swab samples for optimal and consistent DNA yield and quality (Figure 1C) because suboptimal lysis conditions can introduce stochastic bias against gram-positive bacteria because of their thick outer wall. Tests included 1) the Purelink Microbiome DNA Purification Kit (Thermofisher), which is optimized for microorganism lysis, 2) the DNeasy PowerWater Kit (Qiagen), which is optimized for the isolation of genomic DNA from filtered water samples, and 3) phenol-chloroform extraction, which has been shown to produce high DNA yield from environmental samples [29]. The PowerWater kit produced inconsistent yields (results not shown), whereas, the Phenol-Chloroform appoarch produced higher DNA yield, but was countered by pheno carry-over and low DNA purity. The Purelink Microbiome kit consistently produced the highest quality and yield and was subsequently used for the study. Before extraction, frozen filters were crushed in a 2ml Eppendorf tube with sterile 1,000 ml pipette tips. Lysis buffer was added, and bead-beating was performed in a TissueLyser set to full speed for 10 minutes per the manufacturer’s instructions.

### Sequencing

#### Short amplicon

The performance of four amplicon-based 16S-targeting approaches was compared regarding amplification, read quality, taxonomic, and biological conclusions (Figure 1D). Three amplicons designed for short-read Illumina sequencing included 16S variable regions V4 (primers 515F + 806R, hereafter referenced as “Earth”; [30]), V3-4 (primers 341F + 805R, hereafter referenced as “Miseq”; [31], and V1-3 (primers 27F [32] + 534R [33]; hereafter referenced as “27F_534R”; Table 1). One amplicon designed for long-read Pac-Bio sequencing with the Shoreline StrainID kit included 16S, ITS, and 600 bp of the 23S gene [34]; Table 1). The Shoreline Complete StrainID kit uses a patented StrainID primer set.

**Table 1:**
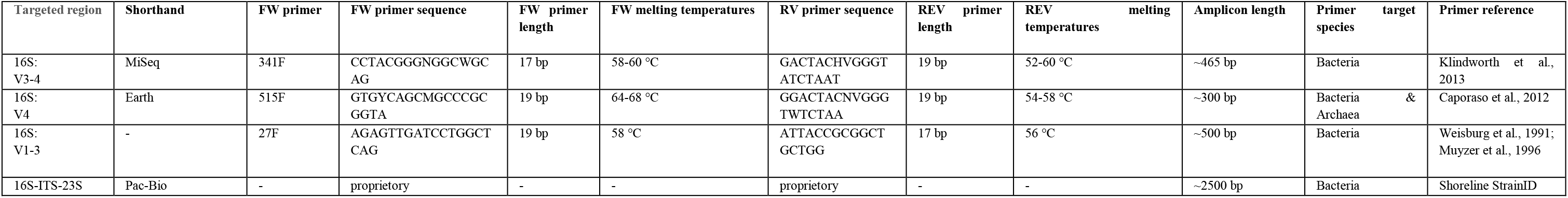
Primers used in this study. FW: forward orientation. REV: reverse orientation. bp: base pairs.

Optimal amplification conditions suitable for all three short amplicons were determined by gradient PCR and reducing cycle number as much as possible. The PCR included 12.5 μl of KAPA HiFi HotStart Ready Mix (Roche, Switzerland), 5 μl of each primer (0.2 μM stock concentration), and 12.5 ng of DNA plus water to a total volume of 25 ul. 14, 16, 18, 20, 22, and 25 PCR cycles were tested at annealing temperatures between 54 and 58°C for all three primer pairs. Based on agarose gel electrophoresis evaluations of amplification success, the following protocol was derived: initial denaturation at 95°C for 3 min, 20 cycles (denaturation at 95°C for 30s, annealing at 55°C for 30s, and extension at 72°C for 20s), and final elongation at 72°C for 5 min. In addition to all samples, four positive controls (Zymobiomics microbial community standard (Zymo Research)) were amplified with this protocol. Samples 19-21 were introduced at this step and are technical PCR-level replicates of sample 2 amplified with Earth primers. Notably, during the sequencing run sample 19 failed to be sequenced.

The preparation of 16S rRNA gene amplicons for the Illumina MiSeq System was designed and performed at the Next Generation Sequencing Platform, University of Bern, according to the “16S Metagenomic Sequencing Library Preparation” protocol (Illumina, art #15044223 Rev. B). The quantity and quality of the cleaned amplicons were assessed using a Thermo Fisher Scientific Qubit 4.0 fluorometer with the Qubit dsDNA HS Assay Kit (Thermo Fisher Scientific, Q32854) and an Agilent Fragment Analyzer (Agilent) with an HS NGS Fragment Kit (Agilent, DNF-474), respectively. Next, the index PCR step was performed as in the protocol except using IDT for Illumina DNA/RNA UD Indexes Set A (Illumina, 20027213), MyFi Mix (BIOLINE, BIO-25050) and the inclusion of a no template control (NTC). The amplicon libraries were assessed for quantity and quality as described above using fluorometry and capillary electrophoresis. The remainder of the protocol was followed, except that the library pool was spiked with 10 % PhiX Control v3 (Illumina, FC-110-3001) to compensate for reduced sequence diversity. Finally, the library was sequenced at 2×300 bp using a MiSeq Reagent Kit v3, 600 cycles (Illumina, MS-102-3003) on the MiSeq sequencing instrument. The run was assessed using Illumina Sequencing Analysis Viewer 2.4.7. We used Illumina bcl2fastq conversion software v2.20 to demultiplex the library samples and convert generated base call files into FASTQ files. All steps beyond the first PCR step were performed at the Next Generation Sequencing Platform, University of Bern. Short-read sequencing, before filtering, resulted in a total of 4,808,910 (27F_534R), 5,306,621 (Earth), and 5,149,263 (MiSeq) reads. Read numbers at all filtering steps are available in Additional File 1. Raw data from Illumina amplicon sequencing are archived in the SRA NBI databank (accession codes to be obtained).

#### Long amplicon

Long amplicon Pac-Bio sequencing was performed at the Next Generation Sequencing Platform, University of Bern. The quantity and quality of the extracted DNA were assessed using a Thermo Fisher Scientific Qubit 4.0 fluorometer with the Qubit dsDNA HS Assay Kit (Thermo Fisher Scientific, Q32854) and an Agilent Femto Pulse system with an Ultra Sensitivity NGS kit (Agilent, FP-1101), respectively. The DNA was then amplified using dual-unique barcoded primers targeting 16S-ITS-23S, using the StrainID kit from Shoreline Biome using strain ID Set Z, Barcodes T1-T16 (Shoreline Biome, STRAIN-Z-SLB). This approach involves a single-step PCR, consisting of primers containing the barcode and target-specific primer, generating amplicons ready for SMRTbell template prep and subsequent sequencing on the Pac-Bio Sequel System. The protocol from input DNA to SMRT sequencing was followed according to the Shoreline Wave for Pac-Bio Technical Manual, following all parameters for the Strain ID workflow. As well as the input DNA of interest, a no template control (NTC), and two community controls (ZymoBIOMICS Microbial Community DNA Standard and ZymoBIOMICS Microbial Community DNA Standard II (Log Distribution) (Zymo Research, D6305 and D6311, respectively) were included. The generated library was SMRT sequenced using a Sequel binding plate 3.0, sequel sequencing plate 3.0 with a 10h movie time on a Pac-Bio Sequel system on their own SMRT cell 1M v3. The library was loaded at 9pM and generated 15 Gb and 284,296 HiFi reads.

Raw data from Pac-Bio amplicon sequencing are from the SRA NBI databank (accession codes between SRR18029933-SRR18029941).

#### Shotgun metagenomics

Illumina shotgun metagenomics sequencing was performed at the Next Generation Sequencing Platform, University of Bern. The extracted DNA was assessed for quantity, purity, and length using a Thermo Fisher Scientific Qubit 4.0 fluorometer with the Qubit dsDNA HS Assay Kit (Thermo Fisher Scientific, Q32854), a DeNovix DS-11 FX spectrophotometer, and an Agilent FEMTO Pulse System with a Genomic DNA 165 kb Kit (Agilent, FP-1002-0275), respectively. Sequencing libraries were made using an Illumina DNA Prep Library Kit (Illumina, 20018705) in combination with IDT for Illumina DNA/RNA UD Indexes Set B, Tagmentation (Illumina, 20027214) according to the Illumina DNA Prep Reference Guide (Illumina, 10000000254 16v09). Six PCR cycles were employed to amplify 30ng of tagemented DNA. Pooled DNA libraries were sequenced paired-end on a NovaSeq 6000 SP Reagent Kit v1.5 (300 cycles; Illumina, 20028400) on an Illumina NovaSeq 6000 instrument. The run produced, on average, 159 million reads/sample. The quality of the sequencing run was assessed using Illumina Sequencing Analysis Viewer (Illumina version 2.4.7) and all base call files were demultiplexed and converted into FASTQ files using Illumina bcl2fastq conversion software v2.20.

Raw data from Illumina shotgun metagenomics sequencing are from the SRA NBI databank (project ID: PRJNA757614).

### Read processing

#### Short amplicon

Illumina short-reads were processed with the DADA2 (Divisive Amplicon Denoising Algorithm 2) [35] pipeline. The DADA2 pipeline includes the inspection of read quality, quality filtering and trimming of reads, dereplication and error rate learning, sample inference for the determination of true sequence variants, merging of reads, construction of sequence table, removal of chimeric reads, and taxonomic assignment. Each primer dataset (Earth, MiSeq, 27F_534R) was first run independently through the DADA2 pipeline, then the Earth and MiSeq datasets were combined and processed identically (“Combined dataset”). The reads were truncated based on the forward and reverse primer read length (Earth forward primer: *trimLeft* – 19; Earth reverse primer: *trimRight* 20; MiSeq forward primer: *trimLeft* – 20; MiSeq reverse primer: *trimRight* 90). For the 27F_534R library, the reads were truncated with *trimLeft* (forward primer: 20, 17) and *trimRight* (Reverse primers: 30, 100). All other *filterAndTrim* parameters were left at the default values. Reads from the individual and combined libraries were merged, while primer pair 27F_534R was concatenated because too many basepairs were trimmed for a successful merging. For the remove bimera denova step, the *minfoldParentOverAbundance* parameter was set to 5 for individual datasets and 4 for the combined dataset. After filtering the datasets retained the following about of reads 3,195,326 (65.8% average) for the 27F_534R dataset, 4,453,843 (83.8%) for the Earth dataset, 4,549,777 (85.5%) for the Earth combined dataset, 4,212,922 (81.7%) for the MiSeq dataset, and 4,216,287 (81.8%) for the MiSeq combined dataset (Additional File 2). Mock community reads were removed from the total amount of reads for each dataset. Additionally, reads from the technical samples (e.g., 20 and 21) were removed for the Earth dataset.

Individual datasets were used to quantify individual primer pair read quality, while the Combined dataset was used to quantify alpha and beta diversity, technical replication reproducibility, and MDS analysis. Sequencing quality was analyzed using the percentage of reads with a Phred score equal to or larger than 30 for each sample type and primer. Microbial taxonomic alpha-diversity (intra-sample) was calculated using Richness and Shannon indices as implemented in the R package phylsoseq [36]. Species beta-diversity (inter-sample) was estimated using the Bray-Curtis dissimilarity metric, while the dissimilarity between groups was visually assessed with multidimensional scaling (MDS) plots.

#### Long amplicon

Pac-Bio Shoreline long reads were demultiplexed without primer trimming, palindromes were removed, and reads with lengths smaller than 200 basepairs were filtered out using the SBAnalyzer software (Shoreline Biome).

#### Shotgun metagenomics

Illumina shotgun metagenomics reads were of high quality, and no filtering was required.

### Taxonomic assignment

#### Short amplicon

Short-read data was assigned to taxonomic units with the SILVA v.138 gene reference database. After DADA2 filtering, the Earth dataset contained 10,941 ASVs, with 196 ASVs assigned to the mock community sample and 3 ASVs assigned to both the mock community and samples. Of the 10,742 ASVs found within the samples, 10,501 were assigned to Bacteria, 14 to Archaea, 57 to Eukaryota, and 2,170 could not be assigned. For the MiSeq dataset, 6,101 ASVs remained, with 22 ASVs assigned to the mock community. Of the 6,079 ASVs found within samples, 6,072 were assigned Bacteria, 2 to Archaea, 2 to Eukaryota, and three could not be assigned. For the combined dataset, 18,075 ASVs remained, with 234 ASVs assigned to the mock community and five ASVs assigned to both the mock community sample and samples. Of the 17,836 ASVs found within the sample, 17,588 were assigned to Bacteria, 16 to Archaea, 58 to Eukaryota, and 174 could not be assigned (Additional File 2).

Sample data were managed using the R package *phyloseq* (v1.30.0) (McMurdie and Holmes, 2013), and plots were generated using the R package *ggplot2* (v.2.2.1) [37].

#### Long amplicon

Long read data were taxonomically assigned with the Athena database v2.2, resulting in 99.3% of reads successfully classified (196,749 reads). An abundance table, a taxonomic classification list for each species, and a list of samples assigned to each read were created (Additional File 3). The initial goal was to compare output after running short- and long-reads through the DADA2 pipeline. However, we obtained low read depths of the samples due to the mock community sample vastly outnumbering the samples during sequencing, which made this approach futile. Therefore, the abundance table was analyzed manually for the spatial distribution of species.

#### Shotgun metagenomics

The raw reads of the metagenomics samples were classified according to their taxonomy using kraken2 [38]. This software classifies reads according to their best matching location in the taxonomic tree. Bracken was used to estimate the species abundance [39], using the taxonomy labels assigned by kraken2 to estimate the number of reads originating from each species present in the sample.

### Data analysis

#### Short amplicon

Read quality was assessed based on the percentage of reads with a Phred score greater than 30 for each primer.

Microbial taxonomic alpha-diversity (intra-sample) was evaluated with the Richness and Shannon indices implemented in the *microbiome* R package [40]. Species beta-diversity (inter-sample) was estimated with Bray-Curtis distances, using the ordinate function in the *phyloseq* package, to understand similarities and differences in community composition independent of primer choice, within-farm compartments, farm identity, and time point in the production cycle. The dissimilarity between samples was assessed by multidimensional scaling (MDS). Statistical variation in microbial alpha-diversity and taxa composition was calculated using a two-way analysis of variance (ANOVA), followed by Post-hoc Tukey tests.

Community composition was analyzed between primers, replicates, sample types, and farms by comparing the relative abundance of the top 9 phyla, all other phyla, and NAs. Furthermore, a PERMANOVA was conducted to determine whether biofilm communities differed significantly between farms.

Finally, ASV enrichments were analyzed with a PERMANOVA non-parametric multivariate test using the adonis function in the R package *vegan* (v.2.5.7) [41] to determine which ASVs were significantly enriched between tank samples of farm A and between farms. The coefficients of the top 20 enriched ASVs were plotted.

All analyses were completed in RStudio 1.4.1717 [42].

#### Long amplicon

The ten most abundant species were identified for each sample type per farm based on the total number of reads after both replicate reads were summed together. Abundance was compiled and plotted for these species to understand spatial and abundance distribution across sample types and farms. Markedly, some replicates have less than ten dots because the top species was only detected in one replicate.

#### Shotgun metagenomics

Phyla with at least 0.5% or more of the total reads were retained to analyze the overall community composition as determined by shotgun sequencing. A Sankey plot using the R network3D v.0.4 package [43] was plotted to compare the community composition across the domains. In addition, relative abundance bar graphs were plotted to quantify community composition variance at the replicate, sample type, and within-farm compartments.

All figures were prepared for publication using Adobe Illustrator 2021.

## Results

### Short amplicon

#### Read quality

Overall read quality was satisfactory, with Earth, MiSeq, and 27F_534R primers producing Phred scores ≥ 30 for 89.3%, 86.6%, and 78.5% of reads, respectively (Figure 2A). However, the lower read quality, combined with the longer amplicon length, led to difficulties during the merging of forward and reverse reads and resulted in overall lower Phred scores for primer pair 27F_534R. Therefore, 27F_534R was not included in downstream processes and analyses.

**Figure 2:**
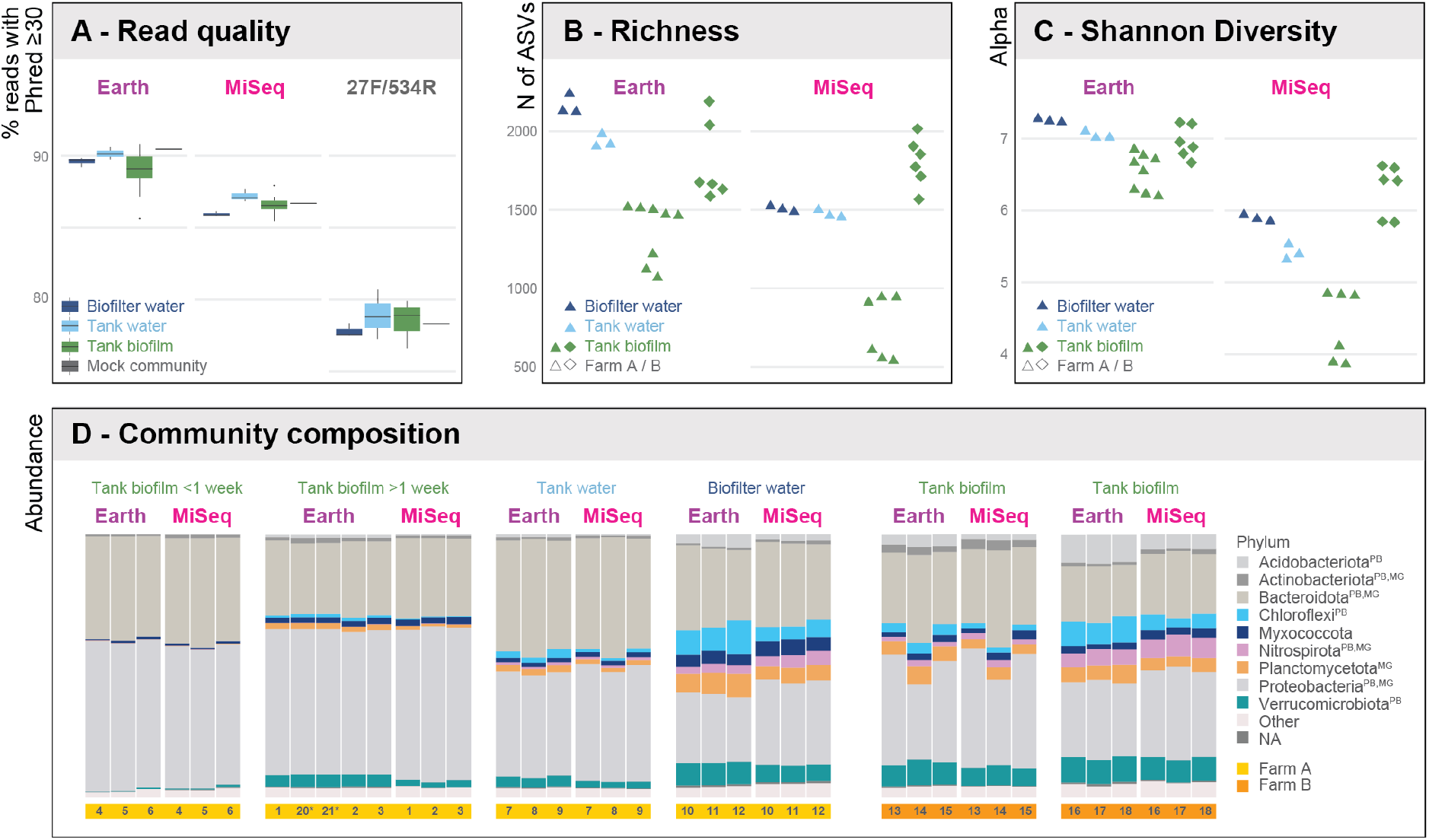
Primer choice affects sequencing and diversity but not community-level results. **A) Read quality.** Earth and MiSeq primers yielded reads with high sequencing quality, whereas primer set 27F/534R yielded lower read quality and was excluded from downstream taxonomic analyses. Between sample types, read quality was comparable for each primer set. **B) Richness.** Earth primers yielded higher alpha diversity based on ASVs across all sample types. In Farm A, biofilter water featured the most ASVs, followed by tank water and finally tank biofilm. Richness in Farm B biofilm was as high (Earth) or higher (MiSeq) than Farm A biofilter water. **C) Diversity.** Samples amplified with Earth primers displayed higher alpha diversity (Shannon index) than samples amplified with MiSeq primers. Patterns are overall similar to richness (panel B). The similar diversity patterns suggest that it would be possible to compare community studies using different primers at the relative scale. **D) Community composition.** The biological and technical replicates (samples 2, 20-21) were highly similar in composition, suggesting that primer selection does not impact spatio-temporal findings at the phyla level and indicates that reproducible results can be obtained with short amplicon sequencing. Indications for succession can be seen in the tank biofilm samples, with increasing complexity from young to older biofilm. Finally, Farm B’s tank biofilm samples resembled Farm A’s biofilter water samples, suggesting that microbial communities with RAS become similar in complexity over time, potentially reaching a stable, mature end. Abbreviations after the phylum name indicate that the phylum was detected in other platform datasets; PB = Pac-Bio, MG = Metagenomics.

#### Taxonomic assignment

Regarding taxonomic assignment, Earth and MiSeq amplicons performed similarly at a higher-level classification (e.g., phylum, order) but diverged at lower-level classification (e.g., genus). Both datasets identified 37 phyla, but the MiSeq dataset assigned 98 more genera at the genus level than the Earth dataset (470 vs. 372) (Additional File 2). Whereas, at the ASV level, the Earth dataset reached higher alpha diversity for the Richness and Shannon indices. In the Earth dataset, ASV richness per sample ranged from 1,067 to 2,240 ASVs, with an average of 1,720.9 ASVs per sample for the combined dataset. In the MiSeq dataset, ASV richness ranged from 437 to 1,948 ASVs, with an average of 1,292.4 ASVs (Figure 2B; Additional File 3). Shannon diversity in the Earth and MiSeq datasets was 6.9 and 5.3, respectively (Figure 2C; Additional File 3).

Between sample types, diversity was highest in biofilter water, followed by tank water and tank biofilm, and impacted by biofilm age. For the Earth dataset, biofilter water had the highest average diversity (Richness: 2,164.3 ASVs, Shannon Diversity: 7.3), followed by tank water (Richness: 1,933.3 ASVs, Shannon Diversity: 7.1). For the MiSeq dataset, biofilm samples from tank two within Farm B had the highest average diversity (Richness: 1810.0 ASVs, Shannon Diversity: 6.1), followed by the biofilm samples from tank one within Farm B (Richness: 1663.3 ASVs, Shannon Diversity: 5.6). Young biofilm samples from Farm A had the lowest diversity in both datasets (Earth: Richness: 1134.3 ASVs, Shannon Diversity: 6.2; MiSeq: Richness: 465.3 ASVs, Shannon Diversity: 4.0) (Additional File 3).

#### Community patterns

Amplicon choice did not affect the composition of the microbial community at higher taxonomic levels. Community composition for distinct sample types, replicates, and the derived spatio-temporal patterns were very similar between the two amplicons (Figures 2D, 3A and B). Subtle biases for/against specific phyla (e.g., *Chloroflexi*, favored by Earth; *Myxococcota* and *Plantomycetota*, favored by MiSeq; Figures 2D) did not affect the inferred overall community structure, which was virtually identical for both amplicons according to MDS analyses (Figures 3A and B).

**Figure 3:**
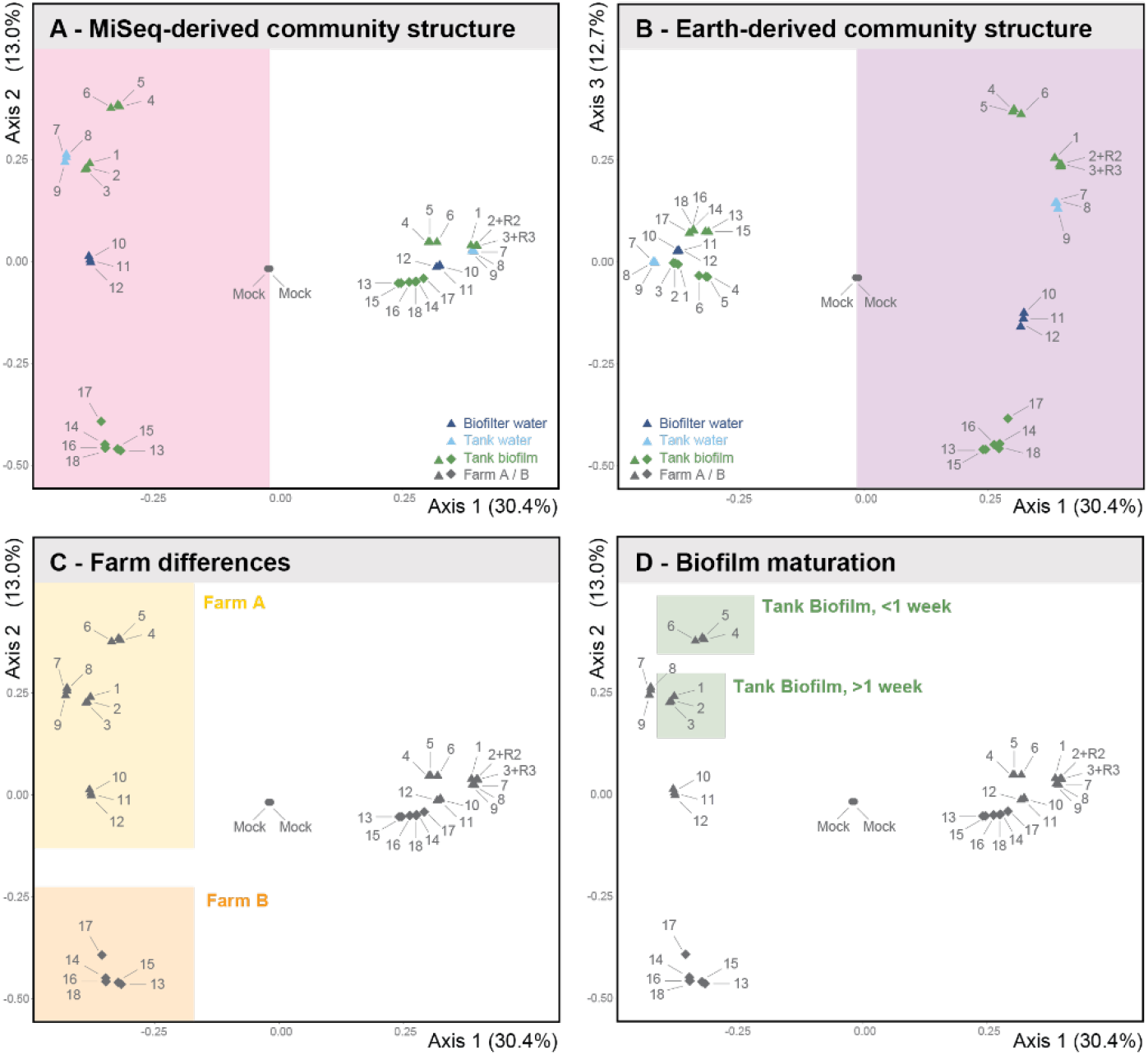
Impact of primers, sample types, farms, and production cycles on community patterns. Multidimensional scaling analysis using the Bray-Curtis distance matrix is visualized in MDS plots at the Phylum level. **A) MiSeqprimers.** Axes 1 and 2 together achieve a clear separation of all samples. Replicates cluster together very closely to the point of overlapping, whereas sample types, compartments, and farms cluster apart. **B) Earth primers.** Earth primers achieve an identical overall pattern but in a distinct area of the morphospace. The mock communities cluster in the exact location independently of primer choice, further stressing the equivalence of primers at this level of analysis. **C) Farms.** Farms separate along Axis 2. Panel C visualizes this for MiSeq primers, but the pattern holds with Earth primers (Panel B). **D) Tank and Time.** Both primers can distinguish the biofilm samples collected in farm A from two tanks of different operational times but within the same circuit, emphasizing that a short-read approach is sufficient to achieve fine-scale resolution at the community level.

Overall community patterns were mostly driven by the variable farm. We found significant differences between farm A and farm B (Earth: Df = 1, F = 9.59, p < 0.00; MiSeq: Df = 1, F = 11.86, p < 0.00; see axis 2 (13.0%) in Figure 3C). This effect was particularly pronounced for tank biofilm (Earth: Df = 1, F = 28.23, p < 0.00; Df = 1, F = 17.25, p < 0.00), with farm B’s biofilm comprising 287 and 356 genera and farm A’s biofilm consisting of 168 and 200 genera for Earth and MiSeq, respectively (Additional File 4).

In addition, distinct sample types featured distinct community compositions (MiSeq: Df = 3, F = 1189, p < 0.000; Earth: Df = 3, F = 250.7, p < 0.000), but same sample types did not necessarily cluster together. For example, farm B’s tank biofilm was more similar to farm A’s biofilter water than farm A’s biofilm samples (Figure 2D). Also, biofilm age drove differences between community richness and dominating genera (Figures 2D and 3D). Biofilm collected one week after tank disinfection vs. biofilm collected several weeks after disinfection featured 113 vs. 153 genera (Earth) or 125 vs. 191 genera (MiSeq) (Figure 2B and C; Additional File 4).

#### Enriched ASVs

The above differences between communities and amplicons were driven by differential enrichment of specific ASVs (Figure 4). When comparing tank water with biofilm for farm A, ASVs affiliated with *Chryseobacteria* were enriched in water and ASVs affiliated with *Rhizobiales, Chtinophagales, Sphareotilus, Ideonella*, and *Flavobacterium* were enriched in the biofilm. This is noteworthy considering the close clustering of these samples in morphospace (Figure 3). When comparing biofilm from farm A and farm B, ASVs enriched in farm A are affiliated with *Rhizobiales, Sphaerotilus, Ideonella*, and *Flavobacteria*, while ASVs affiliated with members of *Aeromonas* and *Flectobacillus* were enriched in farm B (Figure 4).

**Figure 4:**
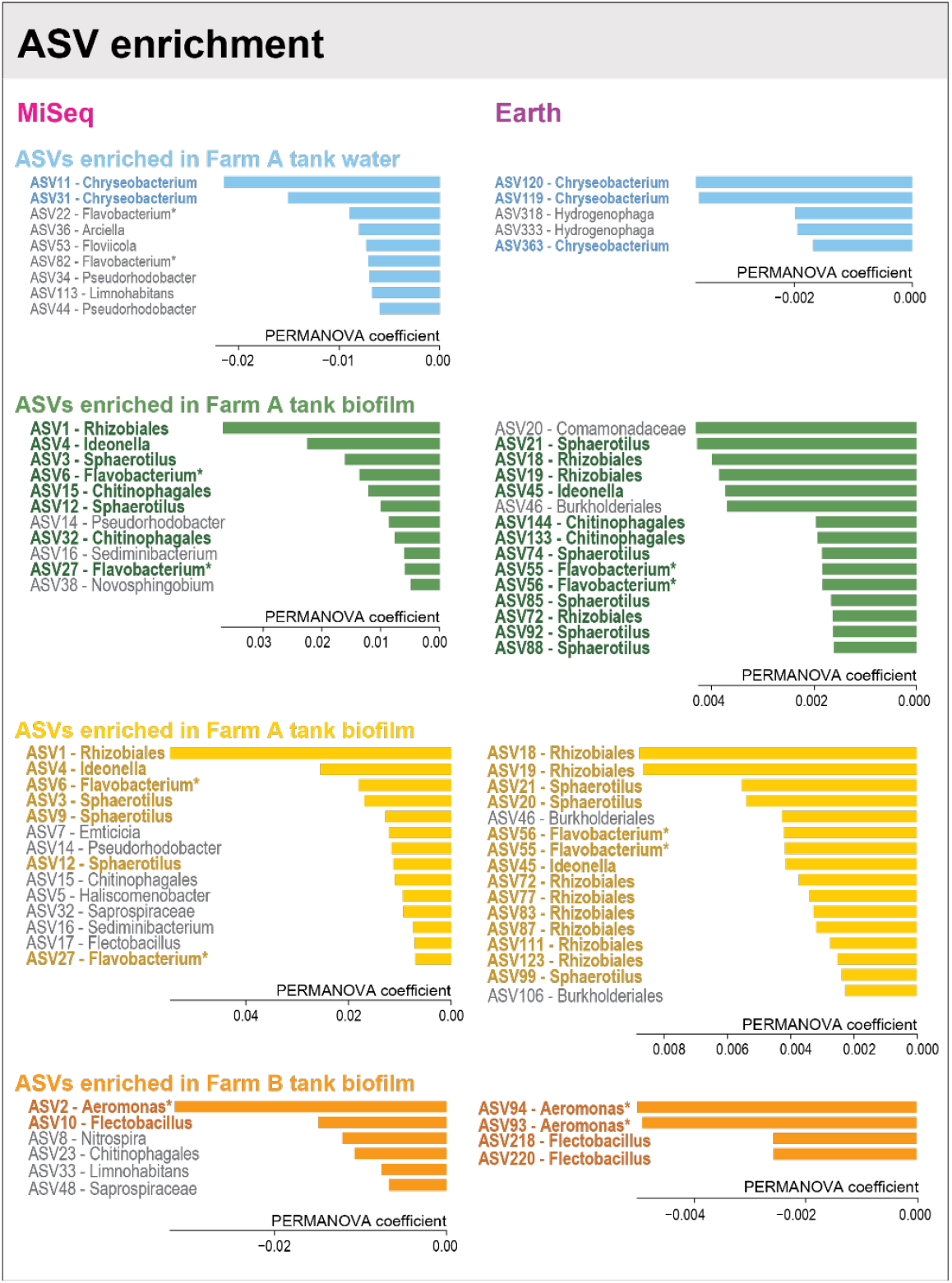
Unique taxonomic units. Permanova coefficients indicate which ASVs are most characteristic for compartments, even though they might not be the most abundant. ASV names are stated in the lowest classified order. ASVs assigned to taxonomic units containing aquaculture pathogens are marked with an asterisk. Primer pair selection starts to emerge at the ASV level, with minor disagreements among the ASVs. For example, Pseudorhodobacter are enriched in farm A using MiSeq primers but not Earth primers. Inversely, Burkholderiales and Comamonadaceae were enriched for Earth, not MiSeq primers. We found that water and biofilm samples from the same farm and circuit differ in enriched ASVs, which is vital for understanding taxa’s diversity and functional services within different sample types. Notably, both primers could correctly assign pathogenic groups to the farm with historical outbreaks of the particular pathogen (i.e., farm A and Flavobacterium; farm B and Aeromonas).

At the level of ASVs, biases are indeed introduced by amplicon choice (Figure 4). For example, ASVs affiliated with *Pseudorhodobacter* are enriched in the MiSeq dataset compared to the Earth dataset in farm A. Conversely, ASVs affiliated with *Burkholderiales* and *Comamonadaceae* are enriched in the Earth dataset. Notably, both Earth and MiSeq datasets were in agreement about the presence of taxonomic groups harboring pathogens (i.e., *Flavobacterium* in farm A and *Aeromonas* in farm B).

### Long amplicon

The low read number obtained from the long-read amplicon approach (a consequence of harsh lysis conditions and over-sequencing of the lower quality sample by higher quality mock community standard) prohibited overall community statistics approaches. Nevertheless, taxonomic conclusions of biological interest could be derived from the 10,041 reads obtained, which resulted in the identification of 204 species (Additional File 5). Similar to the short-read data, species-level data obtained with long-reads emphasize the unique features of farms and, to a lesser extent, compartments (Figure 5). Seventeen of the top enriched species were affiliated with biofilm samples. However, only five were shared between farms, including *Sphaerotilus natans*, a bacterium responsible for bulking, *Streptococcus thermophiles*, a commonly used probiotic bacterium, and *Nitrospira defluvii*, a bacterium that aids nitrification. Twelve species were exclusively detected in farm A, and four were exclusively in farm B. Within farm A, many of the species are detected in at least two compartments, except *Thermomonas sp*. SY21 and *Haliscomenobacter hydrosissis*, which were detected in all compartments. However, species such as *Lysobacter tolerans* and *Paracoccus aminovorans* were found explicitly in farm A’s biofilm. The two water-type samples (biofilter and tank) from the same circuit featured similarities and differences when inspecting the top enriched species, with *Flavobacterium aquatile*, *Propionibacterium freudenreichii*, and *Limnohabitans sp.* 63ED37-2 detected in tank water and *Corynebacterium casei* and *variable*, *Nitrospira defluvii*, and *Brevibacterium yomogidense* detected in biofilter water. Finally, in farm B’s biofilm, *Aeromonas hydrophila*, a common secondary invader known to cause a broad spectrum of infection, was enriched.

**Figure 5:**
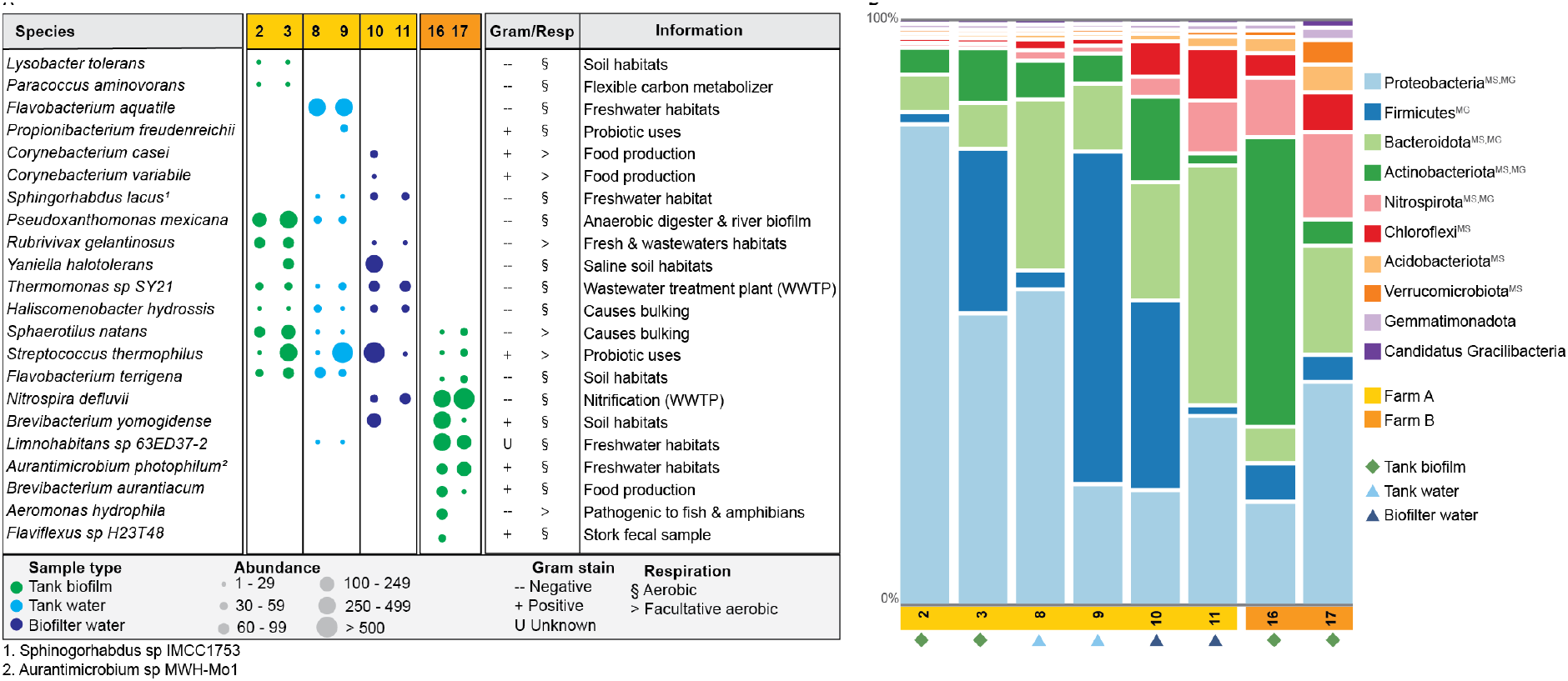
Top 10 species identified with Pac-Bio in each compartment. The top 10 species were identified for each compartment by summing the reads for the replicates and then identifying the 10 most abundant. Updated species names are denoted as footnotes. Green diamonds represent tank biofilm samples, light blue triangles represent tank water, and dark blue triangles represent biofilter water. Pac-Bio data supports that both farms feature distinct communities, with distinct compartments within farm A sharing more species than the same compartment (biofilm) between farms. For example, Haliscomenobacter hydrossis, a species known to cause bulking, is ubiquitous and unique to farm A. Only three species, Sphaerotilus natans, Streptococcus thermophilus, and Flavobacterium terrigena, were detected in both farms, whereas Aeromonas hydrophila was obtained from a tank actively treated for an Aeromonas sp. outbreak. Additional information includes the gram-stain, respiration method, and additional information about the particular species. Abbreviations after the phylum name indicate that the phylum was detected in other platform datasets; MS = MiSeq, MG = Metagenomics.

### Shotgun metagenomics

The shotgun metagenomics data corroborated amplicon findings and extended the picture beyond prokaryotes (75.55%) and included eukaryotes (23.97%), archaea (0.24%), and viruses (0.24%) for farm A samples (Additional file 6). Focusing on phyla with at least 0.5% or more of the total reads, a dataset comprising 96.34% of all reads identified ten phyla. Three-fourths (75.26%) of these reads were assigned to prokaryotic phyla, indicating that competition with eukaryotic reads was not an issue (Figure 6A). Shotgun sequencing agreed with the patterns detected by amplicon sequencing. The top phyla were *Proteobacteria* (54.72% of total reads), *Actinobacteria* (9.47% of total reads), and *Bacteroidetes* (8.05% of total reads) (Fig 6A) for all samples (Fig 6B). The eukaryote phyla comprised *Arthropoda* (10.29%), with insects in fish food as the most likely source; Chordata (8.04%), with the farmed European perch (*Perca flavescens*) as the source; *Ascomycota* (sac fungi); and *Streptophyta* (green algae and plants) (Additional file 6).

**Figure 6:**
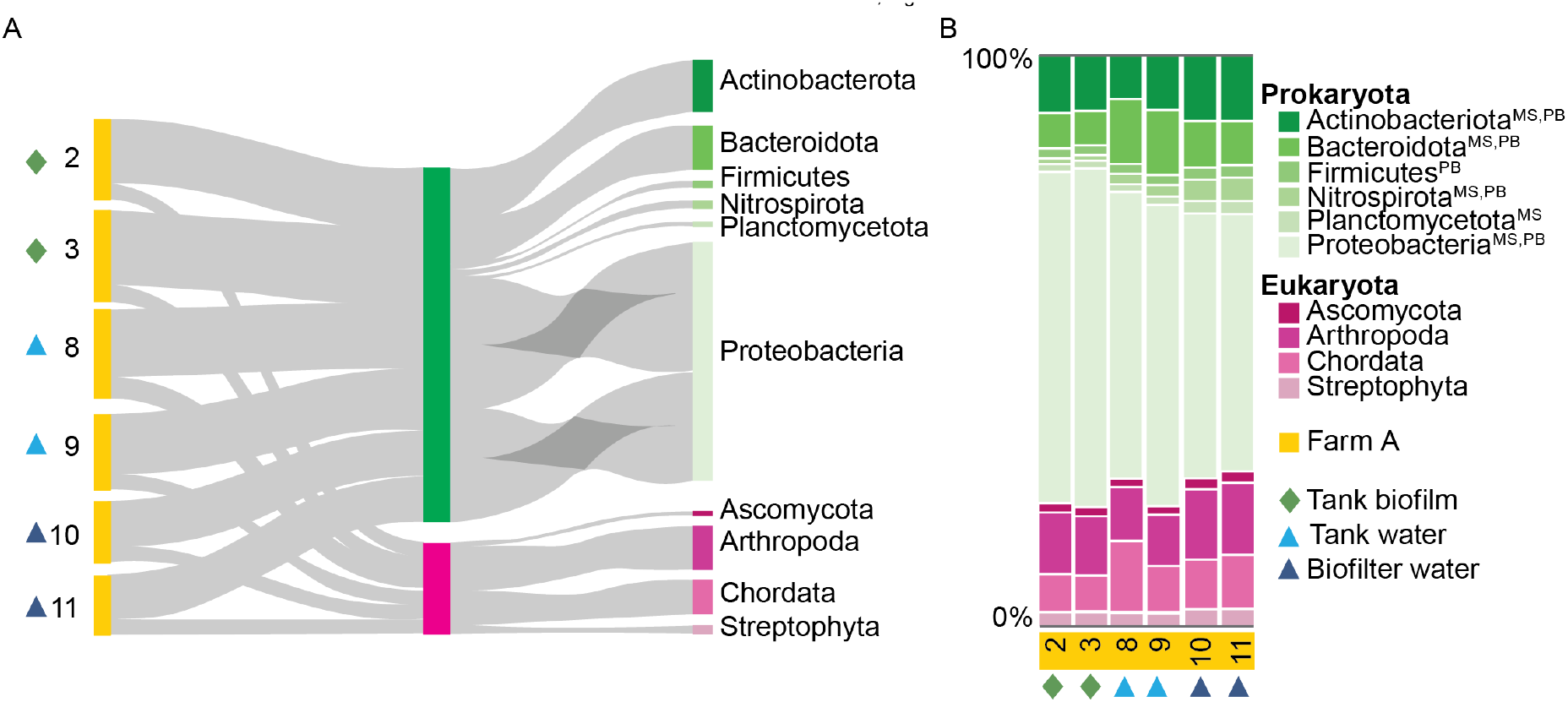
*Phyla with at least* 0.5% or more of the total reads identified with Illumina shotgun metagenomics. A) A Sankey diagram displaying the breakdown between each sample’s prokaryotic and eukaryotic phyla. The prokaryotic phyla are the dominant group for each sample, with Proteobacteria being the most abundant group. Eukaryotic phyla consist of 1) Arthropods likely introduced via the feed, 2) Chordata represented by the farm-raised European perch (Perca fluviatilis), 3) Streptophyta, which consists of green algae and land plants, and 4) Ascomycota, sac fungi, representing the largest phylum of fungi. B) The relative abundance for each sample represents how the community composition changes between replicates, sample types, and within-farm compartments. Overall, the community composition is similar, with some minor differences. For example, the Proteobacteria phyla are less abundant in the biofilter water samples than in tank biofilm samples. Abbreviations after the phylum name indicate that the phylum was detected in other platform datasets; MS = MiSeq, PB = Pac-Bio. Green diamonds represent tank biofilm samples, light blue triangles represent tank water, and dark blue triangles represent biofilter water.

Among lower abundance phyla (0.50 - 0.08% of reads), 16 additional taxa, from a virus group to eukaryotic groups, were detected. The one virus group was *Uroviricota*, a dsDNA-tailed bacteriophages virus. Within the bacteria phyla, four out of the six bacteria were also detected in other datasets. For instance, *Verrucomicrobia*, *Acidobacteriota*, and *Chloroflexi* were also detected in the MiSeq data, and *Gemmatimonadota* was also detected in the Pac-Bio data. In addition, *Euryarchaeota*, a methane-producing archaean, was detected. Lastly, the eukaryotes mainly consisted of non-vertebrate animals such as *Mollusca* (mollusks), *Echinodermata (starfish, sea cucumber and urchins, etc.), Cnidaria (jellyfish, sea anemones, etc.), Nematoda* (roundworms), and *Platyhelminthes* (flatworms). Additionally, *Basidiomycota* (fungus), *Chlorophyta* (green algae), and *Apicomplexa* (protozoan) were detected (Additional file 6).

As expected, pathogenic species were detected at even lower read abundance levels. The top ten pathogenic bacteria, from most abundant to less abundant, included *Flavobacterium psychrophilum* (0.071%)*, Aeromonas veronii* (0.031%)*, A. hydrophila* (0.029%)*, F. branchiophilum* (0.026%)*, F. columnare* (0.015%)*, A. caviae* (0.014%)*, A. salmonicida* (0.005%), *Vibrio vulnificus* (0.004%)*, V. parahaemolyticus* (0.004%), and *A. jandaei* (0.003%) (*Additional file 6*). Noteably, the Pac-Bio data for farm A’s tank water samples also identified *A. hydrophila*, *A. salmonicida*, and *A. veronii*.

## Discussion

Microbial communities are the drivers and determinants of a successful RAS, but their composition and spatio-temporal dynamics are often unknown. Targeted research in RAS is required to shed light on how these communities form, interact and provide services. On the one hand, such knowledge will lead to better management, innovative RAS design, and procedures to manipulate communities. On the other hand, such research will extend our understanding of the rules governing community ecology and evolution beyond controlled lab systems. In this paper, we compare the distinct layers and types of information obtained by distinct methodological approaches from short-read to shotgun metagenomics. We demonstrate that each method can present a cost-effective technique to monitor particular aspects of microbial communities within RAS.

### Primers, Pipelines, and Platforms

Variations in protocols concerning primers and amplification, sequencing platforms, quality filtering, and clustering parameters affect the weight of conclusions in microbial ecology, especially in aquaculture settings, where microbial research lags behind compared to other microbial-based industries. Primer selection for short-read sequencing is potentially the most influential step during a microbial community analysis, as primers directly select for or against specific groups based on the targeted 16S v-region [23], [27], [44]–[46]. Primer bias is, therefore, expected in any study that includes an amplification step, but the main question is how this bias affects the ability to draw sound, reliable, and valid biological conclusions at the desired level of resolution. This is particularly relevant since aquaculture microbiome research widely employs 16S rRNA sequencing as a cost-effective method for surveying microbial communities with minimal starting material requirements [1], [7], [10], [47], [48].

In our study, primer pair 27F_534R underperformed, an unexpected result as this primer pair was successfully used with active sludge collected from a wastewater treatment plant [23]. The shorter MiSeq and Earth amplicons potentially out sequenced 27F_534R at the sequencing step. Therefore, the 27F_534R amplicon, which in theory would offer higher taxonomic resolution due to its increased length [9], [49], could be adequate for RAS samples if not pooled with shorter amplicons. Minor differences in ASV richness between Earth and MiSeq primers did not impact the spatio-temporal patterns and biological conclusion, even though primer bias was detectable at higher taxonomic resolution (Figure 4). Suggesting that community studies can be relatively compared at higher taxonomic levels across studies even though different 16S rRNA primers were used. The significance of these biases is somewhat uncertain, considering that Earth primers have been reported to underestimate the abundance of *Chloroflexi* and *Actinobacteria* in active sludge [23]. In contrast, in our study, Earth primers were biased for *Chloroflexi*, and *Actinobacteria* were similar in their relative abundance between primers (Figure 2D).

In summary, we learned that short-read sequencing is sufficient for understanding spatio-temporal dynamics and community composition at higher taxonomic levels. It is an adequate method for projects that aim to follow spatio-temporal developments at the community level. Because of its low cost, ease of implementation and the availability of well-validated pipelines, 16S rRNA sequencing remains a powerful approach and has potential as a monitoring tool in larger-scale RAS farms that tend to incorporate research and design projects into their annual budgets.

Switching to long-read sequencing approaches is recommended to improve taxonomic resolution [50]–[53] and is desirable in a context where species-specific pathogen identification is relevant. A current drawback is that long-read methods require a large amount of high-quality starting material, thus making them unsuitable for eDNA studies that often have low DNA yield [54] and possibly a high-level of inhibitors. Also, the inclusion of a mock community as recommended to identify biases [25] can compromise sequencing depth. The methodological requirements associated with environmental samples containing gram-positive bacteria, i.e., harsh lysis conditions, compromised our long-read approach that was further impaired when paired with high-quality community standards during sequencing. When aiming for high-quality long-stranded DNA for long-read sequencing, lysis methods and mock standards require additional optimization. We conclude that the taxonomic resolution of the Pac-Bio approach is beneficial in exploring functional services and species identification, especially pathogenic ones, but the approach might not be optimal for a large-scale spatio-temporal study that requires quantitative results.

Shotgun metagenomics does not suffer from primer bias, and genome-wide information, read count and genome size can be used to calculate biogenomic mass - a proxy to determine biomass [55]. Furthermore, species-independent functional profiling based on the presence or absence of genes is another benefit of metagenome data. Finally, RAS microbial ecosystems also harbor archaea [18], fungi [56], [57], and viruses [58], which all interact, compete for resources, and aid or negatively impact the system. Therefore, shotgun metagenomics represents the most powerful -albeit the most expensive - method for quantifying RAS microbial communities. In our study, most reads obtained by shotgun metagenomics were of microbial identity. Furthermore, additional relevant taxa (especially viruses, archaea, and fungi) were detected (Additional file 6), confirming the effectiveness of the approach, and the data patterns mirrored the amplicon data, confirming the validity. The latter further supports the above notion that the impact of primer bias in amplicon approaches is negligible at higher taxonomic levels. Shotgun approaches are, therefore, highly promising and could be further improved by stepping toward an RNA-focused metatranscriptomic approach [59], [60].

Selecting a suited bioinformatics pipeline for analyzing sequencing data is another critical step in microbial studies. Currently, six bioinformatics pipelines are commonly used for 16S rRNA gene amplicon data analysis [61], and all have the potential to introduce bias through sequencing errors [62]. DADA2 is an increasingly used pipeline that shows high sensitivity and can differentiate sequences at single-base resolution, in addition, to clustering with more advanced ASVs [61]. A large body of literature on aquaculture microbiomes works with operational taxonomic units (OTUs), but aquaculture studies using ASVs are on the rise, including studies on host-microbiome interactions [63], microbial dynamics in RAS [10], and microbial dysbiosis during a *Tenacibaculosis* outbreak [64]. We are looking forward to meta-analyses as more ASV-based studies are being published.

Our results support several conclusions on method choice that are probably transferable to other studies. First, primer bias does not compromise higher-level spatio-temporal conclusions of 16S approaches as long as sufficient numbers of high-quality reads are obtained. Importantly, relative differences in community composition between data obtained with different primers can safely be compared, whereas we recommend avoiding comparing absolute statistics of microbial communities analyzed with different primers or lower taxonomic levels. Second, the requirements and challenges of long-read approaches complicate quantitative spatio-temporal community analyses but have value in species-level identification. Lastly, our results agree with other studies on the benefits of hybrid sequencing approaches [65]–[68]. The combination of three different sequencing methods yielded an in-depth overview of spatio-temporal dynamics and species-level information that would otherwise have been difficult to obtain.

### Community composition

The combination of three different sequencing approaches allows for the assessment of potential functional aspects of the analyzed RAS microbial communities. The dominating phyla in both the short-read amplicon (Figure 2D) and the shotgun approach (Figure 6) were *Bacteroidetes* and *Proteobacteria*,which agrees with previous short-read RAS studies (marine RAS: [7], [10], [12], [69]; freshwater RAS: [47]. *Bacteroidetes* contain species that are specialized in the degradation of complex polymers and the cycling of carbon and protein-rich substance [70], [71] and tend to be attached to particles or surfaces [7]. For example, *Flavobacteria*, a class in *Bacteroidetes*, were recently discovered to play a major role in nitrous oxidation-reduction, the final step of denitrification [72]. *Proteobacteria* are a diverse phylum containing nitrifying and denitrifying genera [18], which play a major role in nutrient recycling and remineralization of organic matter [73]–[75], essential steps for the operation of RAS.

A key finding of this study is the impact of the sampling site on results and conclusions. For example, community compositions differed between sampled compartments and sampled time points in the same compartment. Such differences between biofilm and water samples have been reported before, e.g., for a sole RAS [10], a low-through lumpfish farm [8], and an Atlantic salmon RAS [6]. However, all three studies used molecular methods that did not have high enough resolution for species identification, which is essential information for managers when choosing the type of sample to take for diagnostic purposes. In contrast, we used a combination of sequencing data for analyzing the spatial distribution of microbials in RAS.

Within the MiSeq data, the discriminating ASVs for tank water were associated with *Chryseobacterium*, *Flavobacterium*, and *Hydrogenophaga*, whereas biofilm discriminating ASVs were associated with *Rhizobiales*, *Ideonella, Comamonadacea* and *Sphaerotilus*. Importantly, at this level of analysis, conclusions were indeed affected by primer choice, preventing definitive community conclusions. However, two water-associated genera were previously implicated in aspects of fish health, implying that their increased presence in this compartment is of biological significance. *Chryseobacterium* species were reported as emergent fish pathogens across Europe, Asia, and North America. The associated diseases were reported to affect the kidney and/or spleen of cultured rainbow trout (*Oncorhynchus mykiss*), green sturgeon (*Acipenser medirostris*), white sturgeon (*A. tranmontanus*), blue ram cichlid *Mikrogeophagus ramirezi* (a common aquarium fish), and returning fall Chinook salmon (*O. tshawytscha*) from different freshwater systems [76]. Additionally, *Chryseobacterium* species are suspected of playing a role in spoilage [77]–[79] and of being multidrug-resistant [80], which is a danger to both animals and humans. Flavobacterial diseases in fish are caused by multiple bacterial species within the genus *Flavobacterium* and are responsible for devastating losses in wild and farmed fish stock worldwide [81]–[83]. Finally, *Hydrogenophaga* is a group of hydrogen-oxidizing bacteria that perform denitrification services [84]. The identified biofilm-enriched genera have previously been implicated in nutrient-recycling processes. *Rhizobiales* are well-known beneficial partners in plant-microbe interaction and are known to fix nitrogen on legume roots and the stems of some aquatic legumes [85]. However, these farms are not aquaponics, so it is possible that these ASVs are being introduced via the feed and are not a true representation of RAS communities. *Comamonadaceae* also aid in denitrification and removal of phosphorus, thus enhancing water quality for the farmed animals [86]. *Sphaerotilus* is a group often associated with polluted water [87]. Finally, *Ideonella* is a small genus-group composed of four species, with one species, *Ideonella sakaiensis*, capable of degrading PET, a polymer widely used in food containers, bottles, and synthetic fibers [88], and long-read sequencing indicated the presence of this species. Since plastics are used in RAS for biofilter media (e.g., chips, bio-balls), the presence of a potentially plastic-degrading species has implications for replacement and repair cycles.

With the Pac-Bio data, we could not assign ASVs because of low sequencing depth, but it was possible to draw quantitative conclusions between sample types at the species level. For example, within farm A’s biofilm samples, *Lysobacter tolerans* was detected and is a group that produces peptides that can damage the cell walls or membranes of other microbes and are regarded as an untapped source for producing novel antibiotics [89]. In the water samples, *Flavobacterium aquatile*, a known water species typically found in waters containing a high percentage of calcium carbonate – a characteristic of many Swiss waterways [90] and *Propionibacerium freudenreichii* an essential bacteria in the production of Emmental cheese, a Swiss-type cheese [91]. These species were unique to their respective sample type.

Another key notion is the impact of community maturation state on results, especially regarding biofilm communities. Previous studies have detailed the biofilm succession process that entails a non-random process controlled by attachment events, movement, and cellular interactions that induce the non-random spatial organization of biofilms [92]. As biofilms develop, they increase in volume and surface area, creating different gradients of substrates that open niches to varying taxa, such as anaerobic species [93]. This additional complexity increases species richness and functional services, such as degrading organic compounds, cycling of nutrients, or preventing the establishment of pathogenic species. However, biofilms may act as a haven and/or a reservoir for pathogens [94]. For example, *Aeromonas hydrophila* (found in farm B, Additional files 5 and 6) can form thick layers that allow them to evade disinfection or antibiotic treatments [83], [95], while also contributing to the spread of antimicrobial resistance genes [96] - an area we are excited to explore with future shotgun metagenomics data. As a result of these negative impacts, biofilms are often seen as a nuisance and are frequently removed, leaving them in a continuous state of recolonization.

The direct impact of the observed successional processes triggered by repeated biofilm removal on ecological functions and animal health in RAS is unknown. For instance, frequent disruption may potentially open up niches to pathogenic species while preventing the establishment of slower but beneficial species. A study by Rampadarath *et al.* [97] showed that within the first 24 hours of biofilm formation, Proteobacteria microbials were the most dominant, followed by Firmicutes, Bacteroidetes, Chloroflexi, Actinobacteria, and Verrucomicrobia. As previously mentioned, some of the most prominent bacterial fish pathogens are distributed across the phyla Proteobacteria and Bacteroidetes, which could establish early on. Notably, we showed that beneficial *Nitrospira defluvii* are only present in mature samples (farm A: biofilter water and farm B: tank biofilm), suggesting that these species are late colonizers and frequent biofilm removal could prevent their formation, directly impacting the rate of denitrification, potentially threatening animal health. Disentangling the impact of frequent disruption on biofilm communities and the recolonization process is essential in establishing healthy communities after a disruption. Additionally, revealing key bacteria could reduce start-up and operation costs [1], prevent the establishment of pathogens [98], and lead to healthier stock [99].

Finally, community patterns across farms are suggestive of an “island-biogeography” effect, where distinct communities develop in largely isolated habitats. Such effects have been reported in other aquaculture facilities [1], [100]. The long-read data clearly distinguishes farm communities (Figure 5), with *Haliscomenobacter hydrossis* (i.e., causes bulking) [101] and *Streptococcus thermophiles* only being present In farm A. Furthermore, the between-farm biofilm communities only had three species in common: *Sphaerotilus natans*, another bulking species [87], *Streptococcus thermophilus*, and *Flavobacterium terrigena*. In the present case, the conclusion is that farm conditions such as design, management styles, source water, environmental parameters (e.g., temperature, pH), in addition to farmed species, fish feed, and nutrient concentrations [10], [102], combined stochastic assembly processes of dispersal and colonization [103], supersede the continued exchange of microbial communities between farm A and farm B through the delivery of juveniles.

### Disease and Health

Understanding the potential pathogenic risks within a RAS is vital for economic success and animal health. Despite the current growth and optimistic future expectations of the aquaculture sector, the rapid progress of aquaculture has caused unwarranted activities, such as the emergence and spread of pathogens. Aquaculture disease outbreaks can be catastrophic to the industry, causing an estimated worldwide loss of more than US$6.0 billion per anum [104].

Using a shotgun metagenomics approach for farm A’s samples, we detected various pathogenic species that pose a risk to fish health and can ultimately result in massive disease outbreaks (Additional files 5 and 6). *Flavobacterium psychrophilum* (0.092% of total reads, i.e., the causative agent for bacterial coldwater disease) and *Aeromonas veronii* (0.035% of total reads) were the most abundant pathogens detected. Interestingly, they are typically associated with freshwater salmonid fish, such as rainbow trout (*Oncorhynchus mykiss*) and not perch.

### Conclusion

Our results show that microbial communities in RAS are highly dynamic and unique, despite the permanent circulation of water throughout the system. Additionally, management routines create a state of continuous succession and recolonization of these communities, especially biofilm communities. Finally, commonly used 16S primers can detect spatio-temporal development and dynamics between the different RAS compartments, sample types, and farms but cannot prove the resolution required for species or strain identification, critical knowledge for RAS managers.

However, a potential risk is the emergence of a new mutation within these species that would allow for spillover into perch. Functional profiling analyses of the shotgun metagenomics data could shed light on the role functional disease genes play in aquaculture.

Three ubiquitous pathogens known to infect a wide range of firewater fish, including perch, were *Flavobacterium branchiophilum* (i.e., the causative agent of bacterial gill disease), *Aeromonas hydrophila* (i.e., the causative agent of motile aeromonas septicaemia), and *F. columnare* (i.e., the causative agent of columnaris disease). These pathogens were ubiquitous across the system but read counts were the most abundant in tank water. According to farm personnel and in-house confidential diagnostic reports, *F. columnare* and *A. sobria* outbreaks (consisting of 0.002% of the total reads in the shotgun dataset) are the primary pathogens of concern.

Alternative, environmentally-friendly approaches are urgently needed to replace traditional antibiotics for treating bacteria outbreaks in aquaculture. Currently, antibiotics are commonly used to protect or treat against pathogens. Nevertheless, this practice is controversial because of the emergence of antimicrobial resistance genes and the promotion of horizontal gene transfer, in addition to mutagenesis in aquatic bacteria [105]. Furthermore, because of epidemiological connections between humans, animals, and the environment, the increase of antimicrobial resistance genes poses one of the greatest human health and sustainability challenges of the 21^st^ century, according to the World Health Organization [106]. Therefore, developing nonpharmaceutical methods for controlling pathogens is vital for animal and public health. One proposed alternative method is bacteriophage therapy, which uses naturally-occurring bacteriophages to target specific bacteria species or strains of bacteria, such as *Ackermannviridiae* sp. or *Myoviridae* sp., present in both farms (Additional file 6). Unfortunately, phage therapy is still in its infancy, with only a handful of successful phage therapies for the 150 different bacterial pathogens of farmed and wild fish (e.g., *A. hydrophila* in loaches, *F. columnare* in catfish, and *F. psychrophilum* in rainbow trout) [107]. However, as ‘omics technology increases in aquaculture research, this technology could help develop novel mechanisms leading to innovative phage therapies or other pathway-based disruptive measures.

The results presented here contribute to quantifying the microbial community and dynamic and complex interactions in RAS. Further research of microbial communities in aquaculture is necessary to harvest the full power of these micro-but mighty organisms during farm management (e.g., during biofilter startup or disease prevention), to extract basic biological principles (e.g., the link between environmental stressors and microbiome dysbiosis) and to clarify medically relevant interactions (e.g., between host-microbiome-environment interaction and disease development).

## List of Abbreviations

bp: base pairs
FW: forward orientation in regards to primers
RAS: recirculating aquaculture systems
REV: reverse orientation in regards to primers

## Additional Files

Additional file 1: information regarding samples, primers used for amplification, and reads per samples

Additional file 2: information regarding ASVs for each dataset and the assigned taxa

Additional file 3: information regarding the alpha values for each dataset

Additional file 4: information regarding taxa identified during the different maturation stages of biofilm

Additional file 5: information regarding taxa identified with the Pac-Bio sequencing data

Additional file 6: information regarding taxa identified with the Illumina shotgun metagenomics data

Additional files are available upon request from and will accompany the paper upon publication.

## Funding

Swiss National Science Foundation (SNSF) grant: #315230_204838/1 awarded to IAK and CB

## Authors’ contributions

All authors conceived and designed the analysis, CB and IAK acquired funding with the help of JR and AK, JR collected the data, JR and AK performed the data analysis, all authors performed data interpretation, JR wrote the manuscript draft, and all authors edited and reviewed the manuscript.

## Acknowledgments

We want to acknowledge and express our gratitude towards the following people and institutions: Pamela Nicholson and the team of the UniBe NGS Platform for library prep and sequencing, the SIB group of Remy Bruggmann for hosting the Ubelix cluster and providing access to sequencing analysis tools, participating farms for help with sample and data collection and feedback on the manuscript, Loic Marrec for participation in funding acquisition, and FIWI members James Ord and Heike Schmidt-Posthaus for valuable feedback on the manuscript.

## Notes

### Competing Interest Statement

The authors have declared no competing interest.

## References

[1] R. P. Bartelme, M. C. Smith, O. J. Sepulveda-Villet, and R. J. Newton, “Component Microenvironments and System Biogeography Structure Microorganism Distributions in Recirculating Aquaculture and Aquaponic Systems,” mSphere, vol. 4, no. 4, 2019, doi: 10.1128/msphere.00143-19.

[2] J. Dalsgaard, I. Lund, R. Thorarinsdottir, A. Drengstig, K. Arvonen, and P. B. Pedersen, “Farming different species in RAS in Nordic countries: Current status and future perspectives,” Aquac. Eng., vol. 53, pp. 2–13, Mar. 2013, doi: 10.1016/j.aquaeng.2012.11.008.

[3] C. I. M. Martins et al., “New developments in recirculating aquaculture systems in Europe: A perspective on environmental sustainability,” Aquacultural Engineering, vol. 43, no. 3. Elsevier, pp. 83–93, Nov. 01, 2010, doi: 10.1016/j.aquaeng.2010.09.002.

[4] S. Boutin, L. Bernatchez, C. Audet, and N. Derôme, “Network analysis highlights complex interactions between pathogen, host and commensal microbiota,” PLoS One, vol. 8, no. 12, p. e84772, Dec. 2013, doi: 10.1371/journal.pone.0084772.

[5] E. Rurangwa and M. C. J. Verdegem, “Microorganisms in recirculating aquaculture systems and their management,” Rev. Aquac., vol. 7, no. 2, pp. 117–130, Jun. 2015, doi: 10.1111/raq.12057.

[6] I. Bakke et al., “Microbial community dynamics in semi-commercial RAS for production of Atlantic salmon post-smolts at different salinities,” Aquac. Eng., vol. 78, pp. 42–49, Aug. 2017, doi: 10.1016/j.aquaeng.2016.10.002.

[7] I. Rud, J. Kolarevic, A. B. Holan, I. Berget, S. Calabrese, and B. F. Terjesen, “Deep-sequencing of the bacterial microbiota in commercial-scale recirculating and semi-closed aquaculture systems for Atlantic salmon post-smolt production,” Aquac. Eng., vol. 78, pp. 50–62, 2017, doi: 10.1016/j.aquaeng.2016.10.003.

[8] I. Roalkvam, K. Drønen, H. Dahle, and H. I. Wergeland, “Microbial communities in a flow-through fish farm for lumpfish (Cyclopterus lumpus L.) during healthy rearing conditions,” Front. Microbiol., vol. 10, no. JULY, 2019, doi: 10.3389/fmicb.2019.01594.

[9] M. Gołębiewski and A. Tretyn, “Generating amplicon reads for microbial community assessment with next-generation sequencing,” J. Appl. Microbiol., vol. 128, no. 2, pp. 330–354, Feb. 2020, doi: 10.1111/jam.14380.

[10] D. B. Almeida et al., “Microbial community dynamics in a hatchery recirculating aquaculture system (RAS) of sole (Solea senegalensis),” Aquaculture, vol. 539, Jun. 2021, doi: 10.1016/j.aquaculture.2021.736592.

[11] S. Infante-Villamil, R. Huerlimann, and D. R. Jerry, “Microbiome diversity and dysbiosis in aquaculture,” Reviews in Aquaculture, vol. 13, no. 2. John Wiley and Sons Inc, pp. 1077–1096, Mar. 01, 2021, doi: 10.1111/raq.12513.

[12] Y. Ma, X. Du, Y. Liu, T. Zhang, Y. Wang, and S. Zhang, “Characterization of the bacterial communities associated with biofilters in two full-scale recirculating aquaculture systems,” J. Oceanol. Limnol., vol. 39, no. 3, pp. 1143–1150, 2021, doi: 10.1007/s00343-020-0120-8.

[13] S. Moschos, K. A. Kormas, and H. Karayanni, “Prokaryotic diversity in marine and freshwater recirculating aquaculture systems,” Rev. Aquac., Apr. 2022, doi: 10.1111/RAQ.12677.

[14] S. Bagchi et al., “Temporal and spatial stability of ammonia-oxidizing archaea and bacteria in aquarium biofilters,” PLoS One, vol. 9, no. 12, p. e113515, Dec. 2014, doi: 10.1371/journal.pone.0113515.

[15] J. Hüpeden et al., “Relative abundance of Nitrotoga spp. in a biofilter of a cold-freshwater aquaculture plant appears to be stimulated by slightly acidic pH,” Appl. Environ. Microbiol., vol. 82, no. 6, pp. 1838–1845, Mar. 2016, doi: 10.1128/AEM.03163-15.

[16] FAO, The impact of disasters and crises on agriculture and food security: 2021. 2021.

[17] A. Assefa and F. Abunna, “Maintenance of Fish Health in Aquaculture: Review of Epidemiological Approaches for Prevention and Control of Infectious Disease of Fish,” Vet. Med. Int., vol. 2018, 2018, doi: 10.1155/2018/5432497.

[18] H. J. Schreier, N. Mirzoyan, and K. Saito, “Microbial diversity of biological filters in recirculating aquaculture systems,” Current Opinion in Biotechnology, vol. 21, no. 3. Elsevier Current Trends, pp. 318–325, Jun. 01, 2010, doi: 10.1016/j.copbio.2010.03.011.

[19] M. Bentzon-Tilia, E. C. Sonnenschein, and L. Gram, “Monitoring and managing microbes in aquaculture – Towards a sustainable industry,” Microb. Biotechnol., vol. 9, no. 5, pp. 576–584, Sep. 2016, doi: 10.1111/1751-7915.12392.

[20] N. Derome and M. Filteau, “A continuously changing selective context on microbial communities associated with fish, from egg to fork,” Evol. Appl., vol. 13, no. 6, pp. 1298–1319, 2020, doi: 10.1111/eva.13027.

[21] E. U. Yaylacı, “Isolation and characterization of Bacillus spp. from aquaculture cage water and its inhibitory effect against selected Vibrio spp.,” Arch. Microbiol., vol. 204, p. 26, 2022, doi: 10.1007/s00203-021-02657-0.

[22] Z. Huang et al., “Metagenomic analysis shows diverse, distinct bacterial communities in biofilters among different marine recirculating aquaculture systems,” Aquac. Int., vol. 24, no. 5, pp. 1393–1408, Oct. 2016, doi: 10.1007/s10499-016-9997-9.

[23] M. Albertsen, S. M. Karst, A. S. Ziegler, R. H. Kirkegaard, and P. H. Nielsen, “Back to basics - The influence of DNA extraction and primer choice on phylogenetic analysis of activated sludge communities,” PLoS One, vol. 10, no. 7, pp. 31–42, Jul. 2015, doi: 10.1371/journal.pone.0132783.

[24] S.-C. Park and S. Won, “Evaluation of 16S rRNA Databases for Taxonomic Assignments Using a Mock Community,” Genomics Inform., vol. 16, no. 4, p. e24, Dec. 2018, doi: 10.5808/gi.2018.16.4.e24.

[25] J. Pollock, L. Glendinning, T. Wisedchanwet, and M. Watson, “The madness of microbiome: Attempting to find consensus ‘best practice’ for 16S microbiome studies,” Appl. Environ. Microbiol., vol. 84, no. 7, Apr. 2018, doi: 10.1128/AEM.02627-17.

[26] Wasimuddin, K. Schlaeppi, F. Ronchi, S. L. Leib, M. Erb, and A. Ramette, “Evaluation of primer pairs for microbiome profiling from soils to humans within the One Health framework,” Mol. Ecol. Resour., vol. 20, no. 6, pp. 1558–1571, Nov. 2020, doi: 10.1111/1755-0998.13215.

[27] E. Fadeev et al., “Comparison of Two 16S rRNA Primers (V3–V4 and V4–V5) for Studies of Arctic Microbial Communities,” Front. Microbiol., vol. 12, p. 283, Feb. 2021, doi: 10.3389/fmicb.2021.637526.

[28] T. O. Delmont, P. Simonet, and T. M. Vogel, “Describing microbial communities and performing global comparisons in the omic era,” ISME J., vol. 6, no. 9, pp. 1625–1628, 2012, doi: 10.1038/ismej.2012.55.

[29] K. Deiner, J. C. Walser, E. Mächler, and F. Altermatt, “Choice of capture and extraction methods affect detection of freshwater biodiversity from environmental DNA,” Biol. Conserv., vol. 183, pp. 53–63, 2015, doi: 10.1016/j.biocon.2014.11.018.

[30] J. G. Caporaso et al., “Ultra-high-throughput microbial community analysis on the Illumina HiSeq and MiSeq platforms,” ISME J., vol. 6, no. 8, pp. 1621–1624, Aug. 2012, doi: 10.1038/ismej.2012.8.

[31] A. Klindworth et al., “Evaluation of general 16S ribosomal RNA gene PCR primers for classical and next-generation sequencing-based diversity studies,” Nucleic Acids Res., vol. 41, no. 1, pp. e1–e1, Jan. 2013, doi: 10.1093/nar/gks808.

[32] W. G. Weisburg, S. M. Barns, D. A. Pelletier, and D. J. Lane, “16S ribosomal DNA amplification for phylogenetic study,” J. Bacteriol., vol. 173, no. 2, pp. 697–703, 1991, doi: 10.1128/jb.173.2.697-703.1991.

[33] S. H. A. T. Muyzer G., C. Wawer, and Muyzer, G., S. Hottentrager, A. Teske, and C. Wawer, “Denaturing gradient gel electrophoresis of {PCR}-amplified 16S {rDNA}: a new approach to analyze the genetic diversity of mixed,” in Molecular microbial ecology manual, Dordrecht, The Netherlands: Kluwer Academic Publishing, 1996, pp. 1–23.

[34] J. Graf et al., “High-resolution differentiation of enteric bacteria in premature infant fecal microbiomes using a novel rRNA amplicon,” MBio, vol. 12, no. 1, pp. 1–18, 2021, doi: 10.1128/mBio.03656-20.

[35] B. J. Callahan, P. J. McMurdie, M. J. Rosen, A. W. Han, A. J. A. Johnson, and S. P. Holmes, “DADA2: High-resolution sample inference from Illumina amplicon data,” Nat. Methods, vol. 13, no. 7, pp. 581–583, Jun. 2016, doi: 10.1038/nmeth.3869.

[36] P. J. McMurdie and S. Holmes, “Phyloseq: An R Package for Reproducible Interactive Analysis and Graphics of Microbiome Census Data,” PLoS One, vol. 8, no. 4, p. e61217, Apr. 2013, doi: 10.1371/journal.pone.0061217.

[37] P. M. Valero-Mora, “ggplot2: Elegant Graphics for Data Analysis,” Journal of Statistical Software, vol. 35, no. Book Review 1. Springer-Verlag New York, 2010, doi: 10.18637/jss.v035.b01.

[38] D. E. Wood and S. L. Salzberg, “Kraken: ultrafast metagenomic sequence classification using exact alignments,” Genome Biol., vol. 15, no. 3, Mar. 2014, doi: 10.1186/GB-2014-15-3-R46.

[39] J. Lu, F. P. Breitwieser, P. Thielen, and S. L. Salzberg, “Bracken: Estimating species abundance in metagenomics data,” PeerJ Comput. Sci., vol. 2017, no. 1, p. e104, Jan. 2017, doi: 10.7717/PEERJ-CS.104/SUPP-5.

[40] L. Lahti and S. Sudarshan, “Tools for microbiom analysis in R.,” 2012.

[41] J. Oksanen et al., “Package ‘vegan’ Title Community Ecology Package Version 2.5-7,” 2020.

[42] RStudio Team, “RStudio: Integrated development environment for R,” RStudio: Integrated Development Environment for R. Boston, MA, 2019, [Online]. Available: http://www.rstudio.com/.

[43] J. Allaire, C. Gandrud, K. Russell, and C. Yetman, “networkD3: D3 javascript network graphs from r - Google Scholar,” 2017. https://scholar.google.com/scholar?cluster=3312430288369066286&hl=en&oi=scholarr (accessed Jun. 07, 2022).

[44] J. Ghyselinck, S. Pfeiffer, K. Heylen, A. Sessitsch, and P. De Vos, “The effect of primer choice and short read sequences on the outcome of 16S rRNA gene based diversity studies.,” PLoS One, vol. 8, no. 8, p. e71360, Aug. 2013, doi: 10.1371/journal.pone.0071360.

[45] S. Scibetta, L. Schena, A. Abdelfattah, S. Pangallo, and S. O. Cacciola, “Selection and experimental evaluation of universal primers to study the fungal microbiome of higher plants,” Phytobiomes J., vol. 2, no. 4, pp. 225–236, Jan. 2018, doi: 10.1094/PBIOMES-02-18-0009-R.

[46] N. Darwish, J. Shao, L. L. Schreier, and M. Proszkowiec-Weglarz, “Choice of 16S ribosomal RNA primers affects the microbiome analysis in chicken ceca,” Sci. Rep., vol. 11, no. 1, pp. 1–15, Jun. 2021, doi: 10.1038/s41598-021-91387-w.

[47] R. P. Bartelme, S. L. McLellan, and R. J. Newton, “Freshwater recirculating aquaculture system operations drive biofilter bacterial community shifts around a stable nitrifying consortium of ammonia-oxidizing archaea and comammox Nitrospira,” Front. Microbiol., vol. 8, no. JAN, p. 101, Jan. 2017, doi: 10.3389/fmicb.2017.00101.

[48] S. Fu et al., “Horizontal transfer of antibiotic resistance genes within the bacterial communities in aquacultural environment,” Sci. Total Environ., vol. 820, p. 153286, May 2022, doi: 10.1016/J.SCITOTENV.2022.153286.

[49] J. S. Johnson et al., “Evaluation of 16S rRNA gene sequencing for species and strain-level microbiome analysis,” Nat. Commun., vol. 10, no. 1, Dec. 2019, doi: 10.1038/s41467-019-13036-1.

[50] W. Pootakham et al., “High resolution profiling of coral-associated bacterial communities using full-length 16S rRNA sequence data from PacBio SMRT sequencing system,” Sci. Reports 2017 71, vol. 7, no. 1, pp. 1–14, Jun. 2017, doi: 10.1038/s41598-017-03139-4.

[51] T. Klemetsen, N. P. Willassen, and C. R. Karlsen, “Full-length 16S rRNA gene classification of Atlantic salmon bacteria and effects of using different 16S variable regions on community structure analysis,” Microbiologyopen, vol. 8, no. 10, pp. e898–e898, Oct. 2019, doi: 10.1002/mbo3.898.

[52] J. A. Nguinkal et al., “The First Highly Contiguous Genome Assembly of Pikeperch (Sander lucioperca), an Emerging Aquaculture Species in Europe,” Genes 2019, Vol. 10, Page 708, vol. 10, no. 9, p. 708, Sep. 2019, doi: 10.3390/GENES10090708.

[53] L. Tedersoo, M. Albertsen, S. Anslan, and B. Callahan, “Perspectives and Benefits of High-Throughput Long-Read Sequencing in Microbial Ecology,” Appl. Environ. Microbiol., vol. 87, no. 17, pp. 1–19, Aug. 2021, doi: 10.1128/AEM.00626-21.

[54] M. E. Hunter, J. A. Ferrante, G. Meigs-Friend, and A. Ulmer, “Improving eDNA yield and inhibitor reduction through increased water volumes and multi-filter isolation techniques,” Sci. Rep., vol. 9, no. 1, 2019, doi: 10.1038/s41598-019-40977-w.

[55] V. Gaffney et al., “Multi-proxy characterisation of the storegga tsunami and its impact on the early holocene landscapes of the Southern North Sea,” Geosci., vol. 10, no. 7, pp. 1–19, Jul. 2020, doi: 10.3390/GEOSCIENCES10070270.

[56] R. E. Gozlan, W. L. Marshall, O. Lilje, C. N. Jessop, F. H. Gleason, and D. Andreou, “Current ecological understanding of fungal-like pathogens of fish: What lies beneath?,” Front. Microbiol., vol. 5, no. FEB, p. 62, 2014, doi: 10.3389/FMICB.2014.00062/BIBTEX.

[57] E. Dincturk, T. T. Tanrikul, and S. T. Culha, “Fungal and Bacterial Co-Infection of Sea Bass (Dicentrarchus labrax, Linnaeus 1758) in a Recirculating Aquaculture System: Saprolegnia parasitica and Aeromonas hydrophila,” Aquat. Sci. Eng., vol. 33, no. 3, pp. 67–71, 2018, doi: 10.26650/ASE201811.

[58] F. S. Kibenge, “Emerging viruses in aquaculture,” Curr. Opin. Virol., vol. 34, pp. 97–103, Feb. 2019, doi: 10.1016/J.COVIRO.2018.12.008.

[59] M. Shakya, C. C. Lo, and P. S. G. Chain, “Advances and challenges in metatranscriptomic analysis,” Front. Genet., vol. 10, no. SEP, p. 904, Sep. 2019, doi: 10.3389/FGENE.2019.00904/BIBTEX.

[60] C. A. Hempel, N. Wright, J. Harvie, J. S. Hleap, S. J. Adamowicz, and D. Steinke, “Metagenomics vs. total RNA sequencing: most accurate data-processing tools, microbial identification accuracy, and implications for freshwater assessments,” bioRxiv, p. 2022.06.03.494701, Jun. 2022, doi: 10.1101/2022.06.03.494701.

[61] A. Prodan, V. Tremaroli, H. Brolin, A. H. Zwinderman, M. Nieuwdorp, and E. Levin, “Comparing bioinformatic pipelines for microbial 16S rRNA amplicon sequencing,” PLoS One, vol. 15, no. 1, p. e0227434, Jan. 2020, doi: 10.1371/journal.pone.0227434.

[62] R. Bharti and D. G. Grimm, “Current challenges and best-practice protocols for microbiome analysis,” Brief. Bioinform., vol. 22, no. 1, pp. 178–193, Jan. 2021, doi: 10.1093/bib/bbz155.

[63] D. Rosado, M. Pérez-Losada, R. Severino, J. Cable, and R. Xavier, “Characterization of the skin and gill microbiomes of the farmed seabass (Dicentrarchus labrax) and seabream (Sparus aurata),” Aquaculture, vol. 500, pp. 57–64, Feb. 2019, doi: 10.1016/j.aquaculture.2018.09.063.

[64] J. W. Wynne et al., “Microbiome Profiling Reveals a Microbial Dysbiosis During a Natural Outbreak of Tenacibaculosis (Yellow Mouth) in Atlantic Salmon,” Front. Microbiol., vol. 11, p. 586387, 2020, doi: 10.3389/fmicb.2020.586387.

[65] J. M. Lorch et al., “First Detection of Bat White-Nose Syndrome in Western North America,” mSphere, vol. 1, no. 4, Aug. 2016, doi: 10.1128/msphere.00148-16.

[66] G. Fuks et al., “Combining 16S rRNA gene variable regions enables high-resolution microbial community profiling,” Microbiome, vol. 6, no. 1, p. 17, Dec. 2018, doi: 10.1186/s40168-017-0396-x.

[67] D. Bertrand et al., “Hybrid metagenomic assembly enables high-resolution analysis of resistance determinants and mobile elements in human microbiomes,” Nat. Biotechnol. 2019 378, vol. 37, no. 8, pp. 937–944, Jul. 2019, doi: 10.1038/s41587-019-0191-2.

[68] C. L. Brown, I. M. Keenum, D. Dai, L. Zhang, P. J. Vikesland, and A. Pruden, “Critical evaluation of short, long, and hybrid assembly for contextual analysis of antibiotic resistance genes in complex environmental metagenomes,” Sci. Reports 2021 111, vol. 11, no. 1, pp. 1–12, Feb. 2021, doi: 10.1038/s41598-021-83081-8.

[69] M. Brailo et al., “Bacterial community analysis of marine recirculating aquaculture system bioreactors for complete nitrogen removal established from a commercial inoculum,” Aquaculture, vol. 503, pp. 198–206, Mar. 2019, doi: 10.1016/j.aquaculture.2018.12.078.

[70] J. I. Inoue et al., “Distribution and evolution of nitrogen fixation genes in the phylum Bacteroidetes,” Microbes Environ., vol. 30, no. 1, pp. 44–50, 2015, doi: 10.1264/jsme2.ME14142.

[71] X. Shi, K. Kwang Ng, X.-R. Li, and H. Yong Ng, “Investigation of Intertidal Wetland Sediment as a Novel Inoculation Source for Anaerobic Saline Wastewater Treatment,” 2015, doi: 10.1021/acs.est.5b00546.

[72] M. Kuypers, H. Marchant, and B. Kartal, “The microbial nitrogen-cycling network,” Nature Reviews Microbiology, vol. 16, no. 5. pp. 263–276, 2018, doi: 10.1038/nrmicro.2018.9.

[73] D. L. Kirchman, “The ecology of Cytophaga–Flavobacteria in aquatic environments,” FEMS Microbiol. Ecol., vol. 39, no. 2, pp. 91–100, Feb. 2002, doi: 10.1111/J.1574-6941.2002.TB00910.X.

[74] J. P. Zehr, B. D. Jenkins, S. M. Short, and G. F. Steward, “Nitrogenase gene diversity and microbial community structure: A cross-system comparison,” 2003. doi: 10.1046/j.1462-2920.2003.00451.x.

[75] O. V. Tsoy, D. A. Ravcheev, J. Čuklina, and M. S. Gelfand, “Nitrogen fixation and molecular oxygen: Comparative genomic reconstruction of transcription regulation in Alphaproteobacteria,” Front. Microbiol., vol. 7, no. AUG, p. 1343, Aug. 2016, doi: 10.3389/fmicb.2016.01343.

[76] F. De Alexandre Sebastião et al., “Identification of Chryseobacterium spp. isolated from clinically affected fish in California, USA,” Dis. Aquat. Organ., vol. 136, no. 3, pp. 227–234, 2019, doi: 10.3354/DAO03409.

[77] K. Engelbrecht, P. J. Jooste, and B. A. Prior, “Spoilage characteristics of Gram-negative genera and species isolated from Cape marine fish,” S Afr J Food Sci Nutr, vol. 8, pp. 66–71, 1996, Accessed: Jan. 21, 2022. [Online]. Available: https://agris.fao.org/agris-search/search.do?recordID=ZA9600919.

[78] C. Fernández-Álvarez, S. F. González, and Y. Santos, “Development of a SYBR green I real-time PCR assay for specific identification of the fish pathogen Aeromonas salmonicida subspecies salmonicida,” Appl. Microbiol. Biotechnol., vol. 100, no. 24, pp. 10585–10595, Dec. 2016, doi: 10.1007/s00253-016-7929-2.

[79] B. Austin and D. A. Austin, Bacterial fish pathogens: Disease of farmed and wild fish, sixth edition, Sixth. Springer International Publishing, 2016.

[80] J. F. Bernardet, C. Hugo, and B. R. I. T. A. Bruun, “Genus VII. Chryseobacterium Vandamme et al. 1994.” Bergey’s manual of systematic bacteriology,” 2011.

[81] I. Dalsgaard and L. Madsen, “Bacterial pathogens in rainbow trout, Oncorhynchus mykiss (Walbaum), reared at Danish freshwater farms,” J. Fish Dis., vol. 23, no. 3, pp. 199–209, May 2000, doi: 10.1046/j.1365-2761.2000.00242.x.

[82] T. P. Loch and M. Faisal, “Emerging flavobacterial infections in fish: A review,” J. Adv. Res., vol. 6, no. 3, pp. 283–300, May 2015, doi: 10.1016/j.jare.2014.10.009.

[83] W. Cai and C. R. Arias, “Biofilm Formation on Aquaculture Substrates by Selected Bacterial Fish Pathogens,” 2017, doi: 10.1080/08997659.2017.1290711.

[84] M. Lorgen-Ritchie, M. Clarkson, L. Chalmers, J. F. Taylor, H. Migaud, and S. A. M. Martin, “Temporal changes in skin and gill microbiomes of Atlantic salmon in a recirculating aquaculture system – Why do they matter?,” Aquaculture, vol. 558, p. 738352, Sep. 2022, doi: 10.1016/J.AQUACULTURE.2022.738352.

[85] A. Erlacher et al., “Rhizobiales as functional and endosymbiontic members in the lichen symbiosis of Lobaria pulmonaria L,” Front. Microbiol., vol. 6, no. FEB, 2015, doi: 10.3389/fmicb.2015.00053.

[86] Z. Li et al., “Microbial succession in biofilms growing on artificial substratum in subtropical freshwater aquaculture ponds,” FEMS Microbiol. Lett., vol. 364, no. 4, pp. 1–7, 2017, doi: 10.1093/femsle/fnx017.

[87] R. Ferreira et al., “The first sequenced Sphaerotilus natans bacteriophage-characterization and potential to control its filamentous bacterium host,” FEMS Microbiol. Ecol., vol. 97, no. 4, p. 29, Mar. 2021, doi: 10.1093/femsec/fiab029.

[88] G. J. Palm et al., “Structure of the plastic-degrading Ideonella sakaiensis MHETase bound to a substrate,” Nat. Commun., vol. 10, no. 1, pp. 1–10, Apr. 2019, doi: 10.1038/s41467-019-09326-3.

[89] S. Panthee, H. Hamamoto, A. Paudel, and K. Sekimizu, “Lysobacter species: a potential source of novel antibiotics,” Arch. Microbiol., vol. 198, no. 9, pp. 839–845, Nov. 2016, doi: 10.1007/S00203-016-1278-5/FIGURES/4.

[90] B. Müller, J. S. Meyer, and R. Gächter, “Alkalinity regulation in calcium carbonate-buffered lakes,” Limnol. Oceanogr., vol. 61, no. 1, pp. 341–352, Jan. 2016, doi: 10.1002/LNO.10213.

[91] V. de R. Rodovalho, D. L. N. Rodrigues, G. Jan, Y. Le Loir, V. A. de C. Azevedo, and E. Guédon, “<em>Propionibacterium freudenreichii</em>: General Characteristics and Probiotic Traits,” Prebiotics Probiotics - From Food to Heal., Jun. 2021, doi: 10.5772/INTECHOPEN.97560.

[92] M. Grinberg, T. Orevi, and N. Kashtan, “Bacterial surface colonization, preferential attachment and fitness under periodic stress,” PLOS Comput. Biol., vol. 15, no. 3, p. e1006815, Mar. 2019, doi: 10.1371/JOURNAL.PCBI.1006815.

[93] C. Suarez, M. Piculell, O. Modin, S. Langenheder, F. Persson, and M. Hermansson, “Thickness determines microbial community structure and function in nitrifying biofilms via deterministic assembly,” Sci. Rep., vol. 9, no. 1, pp. 1–10, 2019, doi: 10.1038/s41598-019-41542-1.

[94] H. C. Flemming, J. Wingender, U. Szewzyk, P. Steinberg, S. A. Rice, and S. Kjelleberg, “Biofilms: an emergent form of bacterial life,” Nat. Rev. Microbiol. 2016 149, vol. 14, no. 9, pp. 563–575, Aug. 2016, doi: 10.1038/nrmicro.2016.94.

[95] W. Cai, L. De La Fuente, and C. R. Arias, “Biofilm formation by the fish pathogen flavobacterium columnare: Development and parameters affecting surface attachment,” Appl. Environ. Microbiol., vol. 79, no. 18, pp. 5633–5642, 2013, doi: 10.1128/AEM.01192-13.

[96] D. Stratev and O. A. Odeyemi, “Antimicrobial resistance of Aeromonas hydrophila isolated from different food sources: A mini-review,” J. Infect. Public Health, vol. 9, no. 5, pp. 535–544, Sep. 2016, doi: 10.1016/J.JIPH.2015.10.006.

[97] S. Rampadarath, K. Bandhoa, D. Puchooa, R. Jeewon, and S. Bal, “Early bacterial biofilm colonizers in the coastal waters of Mauritius,” Electron. J. Biotechnol., vol. 29, pp. 13–21, Sep. 2017, doi: 10.1016/J.EJBT.2017.06.006.

[98] N. Derome, J. Gauthier, S. Boutin, and M. Llewellyn, “Bacterial Opportunistic Pathogens of Fish,” Springer, Cham, 2016, pp. 81–108.

[99] K. J. K. Attramadal, T. M. H. Truong, I. Bakke, J. Skjermo, Y. Olsen, and O. Vadstein, “RAS and microbial maturation as tools for K-selection of microbial communities improve survival in cod larvae,” Aquaculture, vol. 432, pp. 483–490, Aug. 2014, doi: 10.1016/j.aquaculture.2014.05.052.

[100] V. Schmidt, L. Amaral-Zettler, J. Davidson, S. Summerfelt, and C. Good, “Influence of fishmeal-free diets on microbial communities in atlantic salmon (Salmo Salar) recirculation aquaculture systems,” Appl. Environ. Microbiol., vol. 82, no. 15, pp. 4470–4481, 2016, doi: 10.1128/AEM.00902-16/SUPPL_FILE/ZAM999117249SO1.PDF.

[101] C. Cao and I. Lou, “Analysis of environmental variables on population dynamic change of Haliscomenobacter hydrossis, the bulking causative filament in Macau wastewater treatment plant,” Desalin. Water Treat., vol. 57, no. 16, pp. 7182–7195, Apr. 2016, doi: 10.1080/19443994.2015.1014857.

[102] L. N. Duarte, F. J. R. C. Coelho, D. F. R. Cleary, D. Bonifácio, P. Martins, and N. C. M. Gomes, “Bacterial and microeukaryotic plankton communities in a semi-intensive aquaculture system of sea bass (Dicentrarchus labrax): A seasonal survey,” Aquaculture, vol. 503, pp. 59–69, Mar. 2019, doi: 10.1016/j.aquaculture.2018.12.066.

[103] J. Zhou, “Stochastic Community Assembly: Does It,” Microbiol. Mol. Biol. Rev., vol. 81, no. 4, pp. 1–32, 2017.

[104] S. S. Mishra, R. Das, and P. Swain, “Status of Fish Diseases in Aquaculture and assessment of economic loss due to disease,” 2019, pp. 183–198.

[105] T. Nogueira and A. Botelho, “No Title,” vol. 10, no. 7, Jul. 2021, Accessed: Jun. 10, 2022. [Online]. Available: /pmc/articles/PMC8300701/.

[106] WHO, “Antimicrobial resistance,” Antimicrobial resistance, 2021. https://www.who.int/news-room/fact-sheets/detail/antimicrobial-resistance (accessed Jun. 20, 2022).

[107] G. P. Richards, “Bacteriophage remediation of bacterial pathogens in aquaculture: a review of the technology,” Bacteriophage, vol. 4, no. 4, p. e975540, Dec. 2014, doi: 10.4161/21597081.2014.975540.

